# Effects of axon branching and asymmetry between the branches on transport, mean age, and age density distributions of mitochondria in neurons: a computational study

**DOI:** 10.1101/2022.04.25.489429

**Authors:** Ivan A. Kuznetsov, Andrey V. Kuznetsov

## Abstract

We report a computational study of mitochondria transport in a branched axon with two branches of different sizes. For comparison, we also investigate mitochondria transport in an axon with symmetric branches and in a straight (unbranched) axon. The interest in understanding mitochondria transport in branched axons is motivated by the large size of arbors of dopaminergic neurons, which die in Parkinson’s disease. Since the failure of energy supply of multiple demand sites located in various axonal branches may be a possible reason for the death of these neurons, we were interested in investigating how branching affects mitochondria transport. Besides investigating mitochondria fluxes between the demand sites and mitochondria concentrations, we also studied how the mean age of mitochondria and mitochondria age densities depend on the distance from the soma. We established that if the axon splits into two branches of unequal length, the mean ages of mitochondria and age density distributions in the demand sites are affected by how the mitochondria flux splits at the branching junction (what portion of mitochondria enter the shorter branch and what portion enter the longer branch). However, if the axon splits into two branches of equal length, the mean ages and age densities of mitochondria are independent of how the mitochondria flux splits at the branching junction. This even holds for the case when all mitochondria enter one branch, which is equivalent to a straight axon. Because the mitochondrial membrane potential (which many researchers view as a proxy for mitochondrial health) decreases with mitochondria age, the independence of mitochondria age on whether the axon is symmetrically branched or straight (providing the two axons are of the same length), and on how the mitochondria flux splits at the branching junction, may explain how dopaminergic neurons can sustain very large arbors and still maintain mitochondrial health across branch extremities.

## 1. Introduction

Dopaminergic (DA) neurons in the substantia nigra pars compacta, which die during Parkinson’s disease (PD) [1], are characterized by large, unmyelinated axonal arbors. The morphology of these arbors is extremely complex. The axon of a DA neuron splits at multiple branching junctions into numerous branches. In humans, a single DA axon can form more than a million synapses [2]. Maintaining such a large arbor requires a lot of energy, and problems with energy supply may explain the reason for the death of DA neurons in PD [3].

Mitochondria are continuously transported in axons [4], and probably one of the main reasons for this transport (unless we consider an axon with a growing neuron) is maintaining mitochondrial health. One important question relevant to DA neurons is how mitochondria traffic is maintained in axons with multiple branches. A second is how mitochondria age is affected by the branched paths inherent to mitochondria transport in such axons [5]. Mitochondria are too large to exhibit any significant diffusivity: the representative length of mitochondria is between 0.5 and 10 μm [6]. Therefore, mitochondria are transported by molecular motors; long-range anterograde motion along microtubules (MTs) is due to kinesin-1 (and possibly by kinesin-3) motors and retrograde motion along MTs is due to cytoplasmic dynein [7,8]. Mitochondria can also stop their motion and become anchored at various energy demand sites (hereafter demand sites) in the axon [9-11].

Like anything else in neurons, mitochondria do not last forever; damage by reactive oxygen species may be one of the reasons for this [12]. Therefore, an entire mitochondrion, or mitochondrial proteins, need to be periodically replaced [13]. Since the death of DA neurons in PD may be caused by defects in mitochondria transport, investigating the mean age of mitochondria in an axon, and how the mean age is affected by axon branching, is relevant to understanding the fundamentals of PD.

Delivery of motor-cargo complexes to branched axons was investigated using mean-field equations in [14]. Mitochondrial health can be treated as a proxy for mitochondrial membrane potential [15]. Ref. [16] developed models that treat mitochondrial health as a decaying component; they reported simulations of mitochondrial health in straight and branched axons. In this paper, we develop a different, compartment-based model which simulates transport of mitochondria between various demand sites. Each demand site is represented by three compartments that contain populations of anterograde, stationary, and retrograde mitochondria, respectively.

We simulated mitochondria transport in an axon that splits into two unequal branches (Fig. 1a). We investigated the following questions: (i) what happens with mitochondria concentrations when an axon splits into two branches; (ii) what happens with mitochondria age when an axon splits into two branches; (iii) how the asymmetry in branches affects the mean age and the age density distribution of mitochondria in the branches and the segment of the axon before the branching junction; (iv) how the splitting of the flux at the branching junction affects the mean age and age density distribution of mitochondria.

**Fig. 1.**
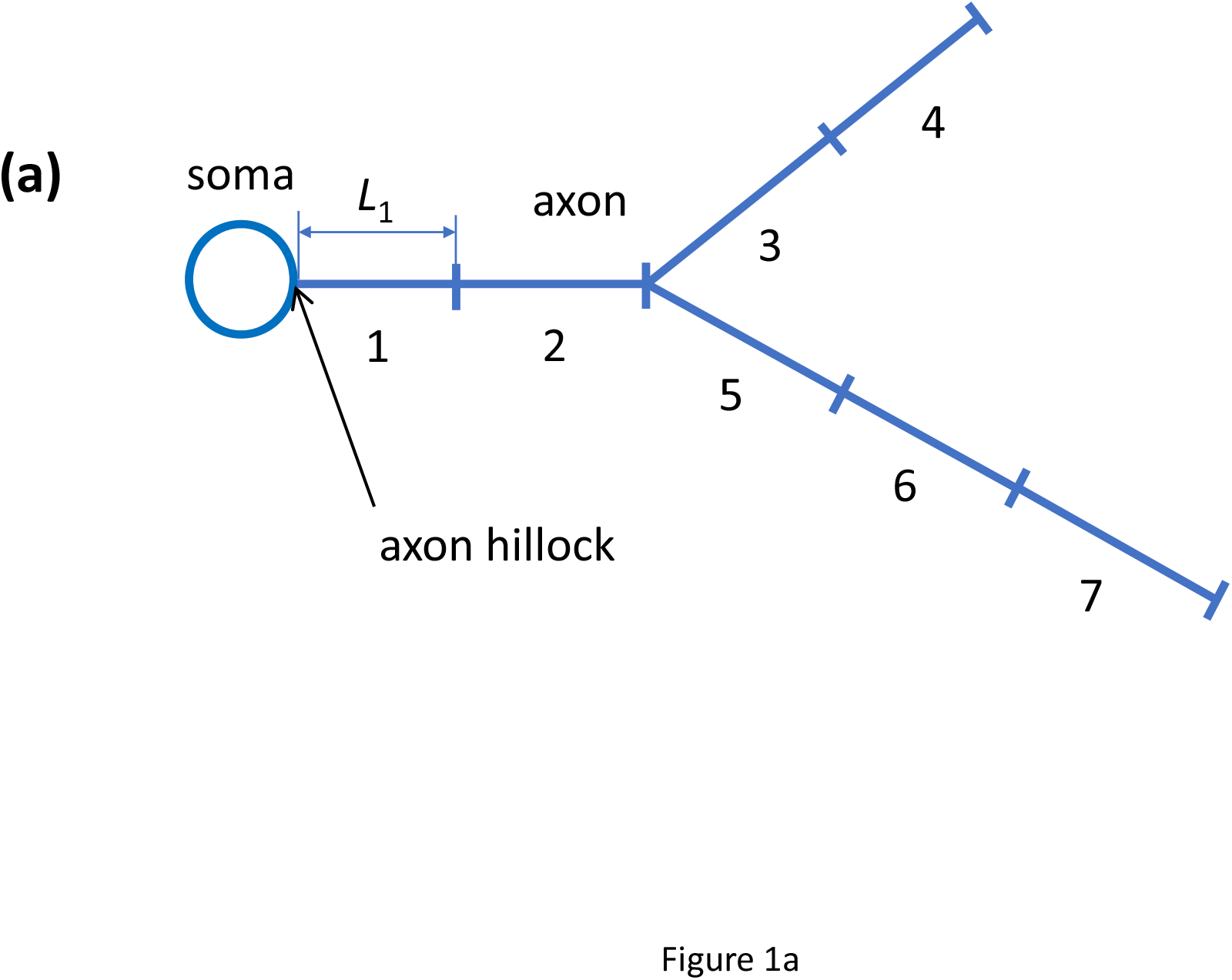
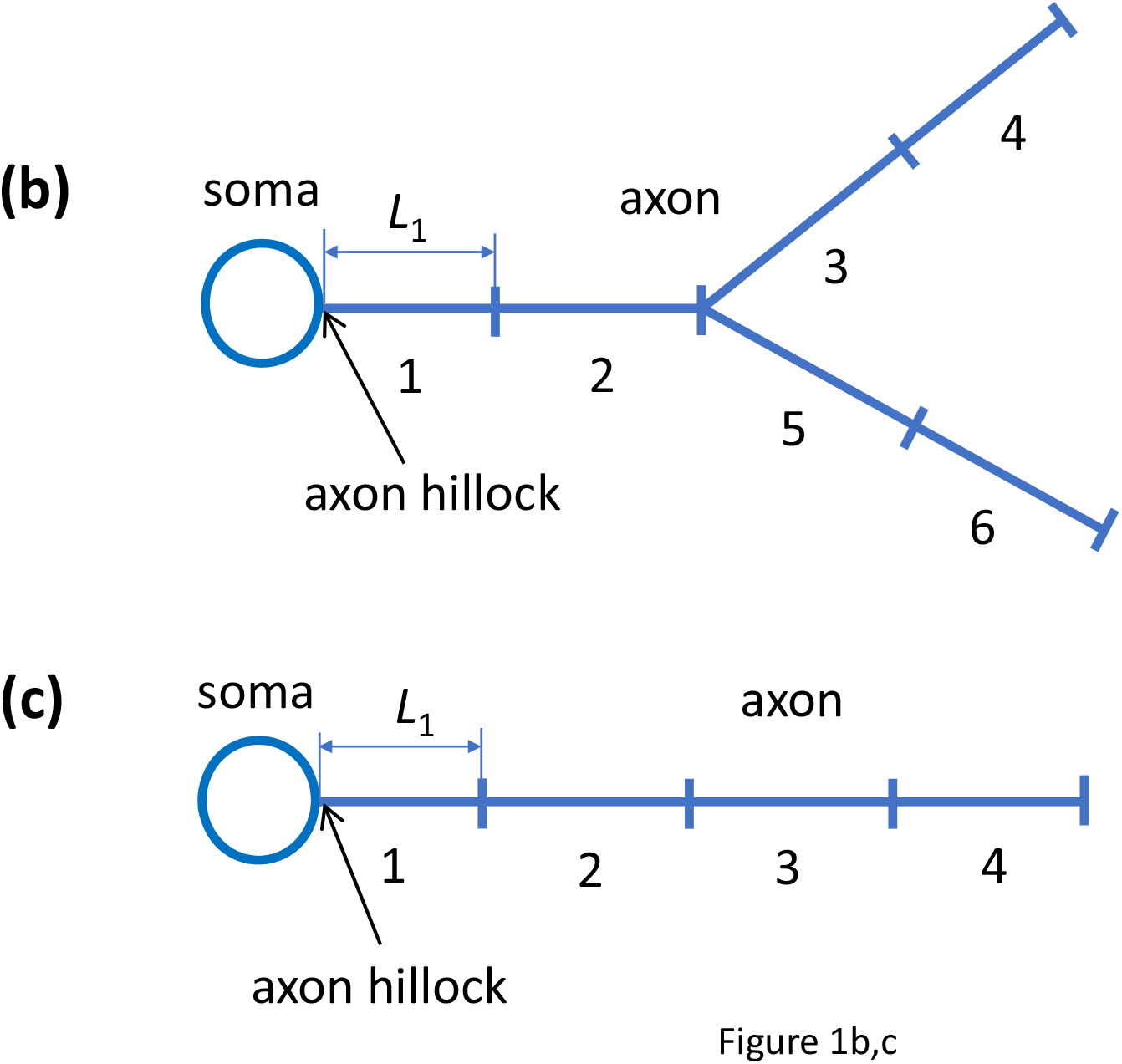
A schematic diagram of the compartmental representation of a neuron with an asymmetric branched axon (a), with a symmetric branched axon (b), and a neuron with a non-branched (straight) axon (c). The length of the compartment representing the most proximal demand site is shown; all other compartments are assumed to be of the same length.

To answer these questions, we compared mitochondria transport in a branched axon with one shorter and one longer branch (Fig. 1a) with that in an axon with two identical branches (Fig. 1b) and in a non-branched, straight axon (Fig. 1c). The model was obtained by extending the model of mitochondria transport reported in [17]. The mean age and age density distributions of mitochondria in various compartments were computed utilizing the methods developed in [18] and [19].

## 2. Materials and models

### 2.1. Equations expressing conservation of the total length of mitochondria in various compartments for an axon with two asymmetric branches (Figs. 1a and 2a)

Mitochondria transport in the axon is simulated by a multi-compartment model [20-22]. The reason for choosing this model is our interest in simulating the mean age and age density distributions of mitochondria in various locations in the axon and axonal branches. We thus utilize the methodology developed in [18] and [19] that allows the calculation of the age of particles transported in compartmental systems.

Since our goal is to investigate the effect of axon branches of unequal length on mitochondria age distributions in the axon, we represented the axon by seven demand sites: two in the segment before the branching junction, two representing the short branch, and three representing the long branch (Fig. 1a). Each demand site is represented by three compartments: one with anterograde mitochondria, one with stationary mitochondria, and one with retrograde mitochondria (Fig. 2a). Mitochondria can transition between the compartments. Also, anterograde mitochondria can move to the next demand site and retrograde mitochondria can return to the previous demand site. Mitochondria continuously undergo fusion and fission. Since tracking mitochondria with different lengths is difficult by a model that utilizes the mean-field equations, we lumped the lengths of all mitochondria contained within the same compartment. Our model simulates the total length of mitochondria in a compartment, which is a conserved property. Since axons are elongated in one direction, we evaluate mitochondria concentrations by the total length of mitochondria residing in a certain kinetic state (anterograde, stationary, or retrograde) contained in a unit length of the axon. The unit for mitochondria concentration thus is (µm of mitochondria)/µm. For the axon displayed in Fig. 1a, whose compartmental representation is shown in Fig. 2a, the model simulates 27 mitochondria concentrations in 27 different compartments. As in any compartmental system, the only independent variable in our model is time, *t*. We summarized the dependent variables for our model in Table S1.

**Fig. 2.**
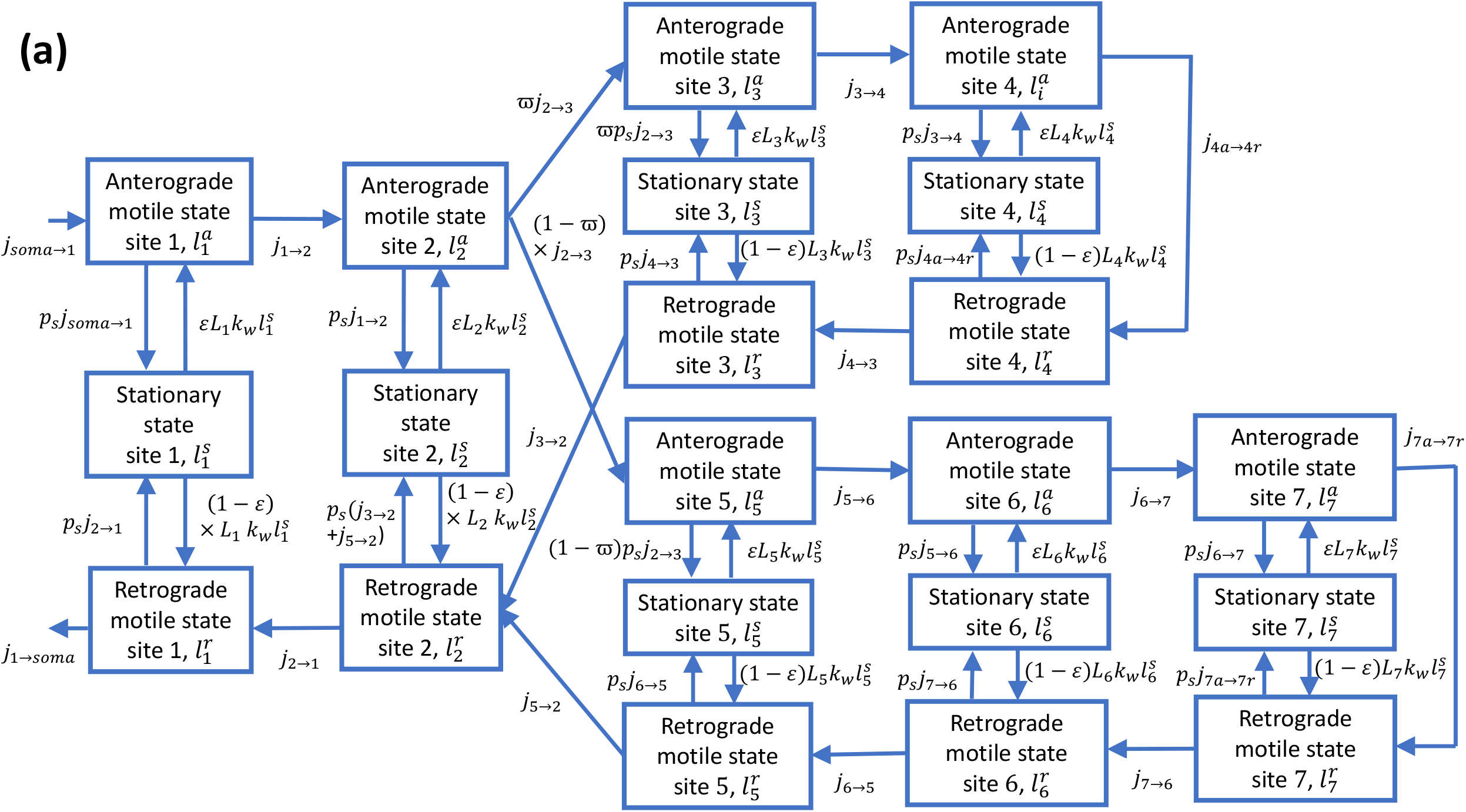
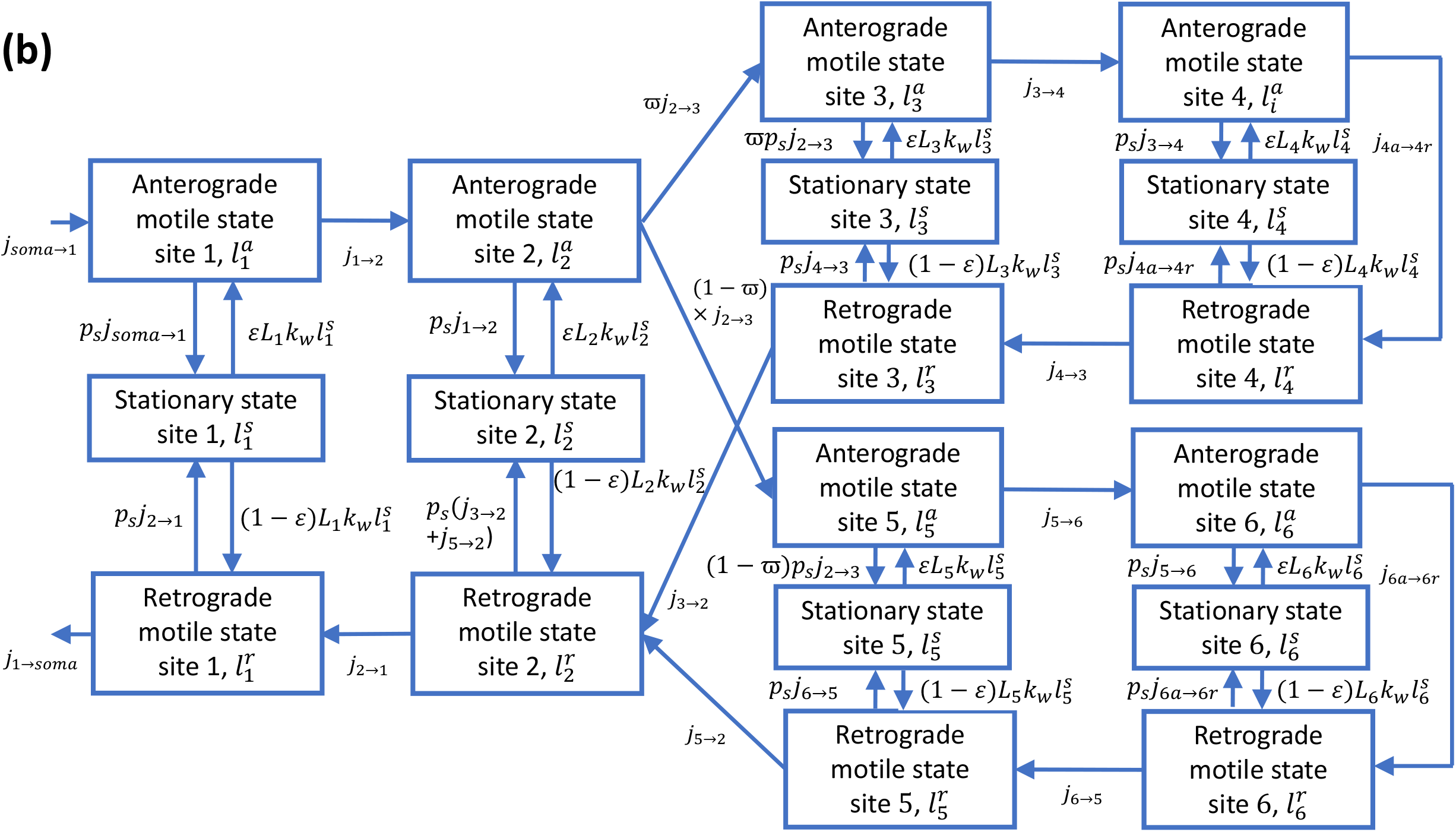
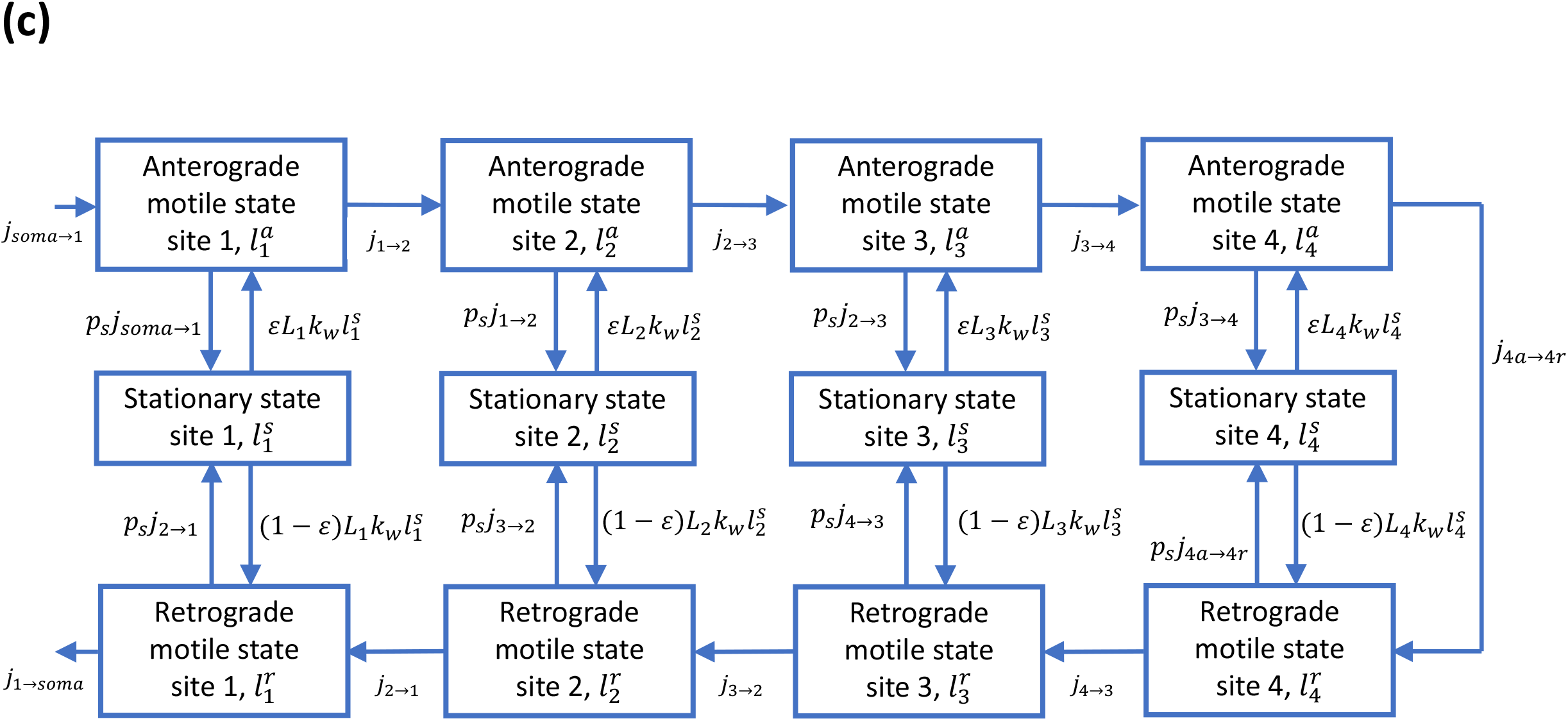
A kinetic diagram showing a compartmental model representation of an asymmetric branched (a), symmetric branched (b), and non-branched (straight) (c) axon. Compartments represent demand sites in the terminal. Each demand site is represented by three compartments: those with anterograde, stationary, and retrograde mitochondria. The mitochondrial exchange between the transiting (anterograde and retrograde) states in adjacent demand sites is depicted by arrows. We assumed that mitochondria entering the terminal have zero age. The capture of mobile mitochondria into the stationary states and their release from the stationary states are also shown by arrows.

The conservation of the total length of stationary mitochondria in the most proximal demand site (# 1) is given by the following equation (Fig. 2a):

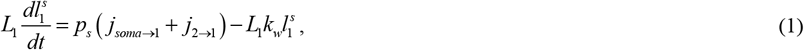

where 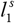 is the concentration of stationary mitochondria in the most proximal compartment and *L*_1_ is the length of the axonal compartments around the first demand site. For all axons depicted in Fig. 1, we assumed that *L*_1_ = *L*_*ax*_ /4. If the axon is asymmetrically branched, we interpreted *L*_*ax*_ as the distance from the hillock to the tip of the shorter branch. This was done to keep the length of compartments representing demand sites the same for all cases depicted in Figs. 1a, 1b, and 1c. All other axonal compartments (*L*_2_, *L*_3_, etc.) were assumed to be of the same length. Also, *p*_*s*_ is the probability for a motile mitochondrion to transition to a stationary state when it passes a demand site, *k*_*w*_ is the kinetic constant characterizing the rate at which stationary mitochondria reenter the motile states (anterograde and retrograde), and *j*_*soma*→1_ is the rate of mitochondria production in the soma (all mitochondria produced in the soma are assumed to enter the axon). All other fluxes between the compartments containing motile mitochondria, *j*_*a*→*b*_, are given by Eqs. (22)-(35) below.

The conservation of the total length of anterograde mitochondria in the most proximal demand site (# 1) is given by the following equation (Fig. 2a):

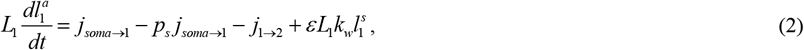

where 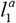 is the concentration of anterograde mitochondria in the most proximal compartment and *ε* is a model constant that simulates what portion of mitochondria that are released from the stationary state join the anterograde pool; the remaining portion, (1− *ε*), join the retrograde pool.

The conservation of the total length of retrograde mitochondria in the most proximal demand site (# 1) is given by the following equation (Fig. 2a):

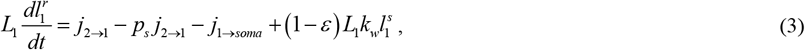

where 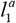 is the concentration of retrograde mitochondria in the most proximal compartment.

Stating the conservation of the total length of mitochondria in the compartments contacting stationary, anterograde, and retrograde mitochondria around other demand sites (#2,…,7) gives the following equations:

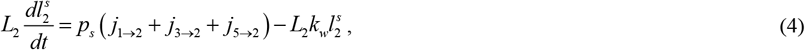

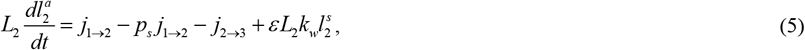

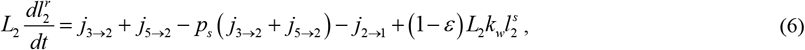

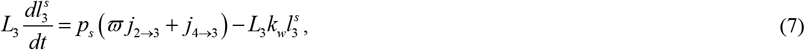

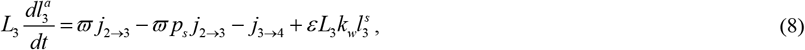

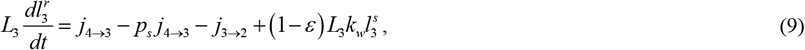

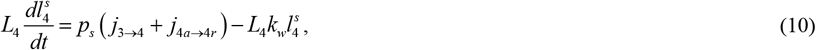

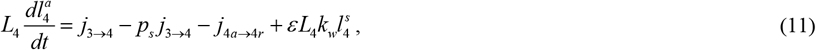

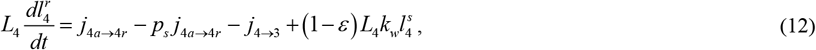

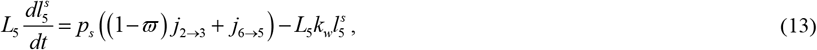

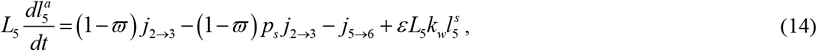

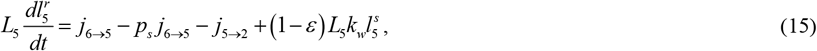

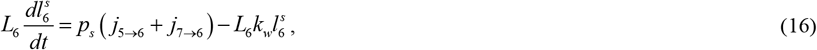

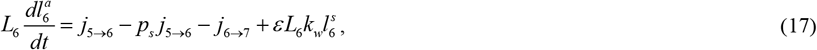

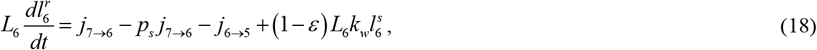

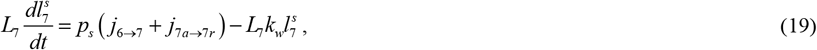

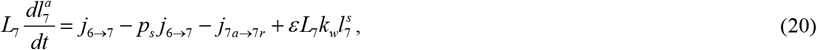

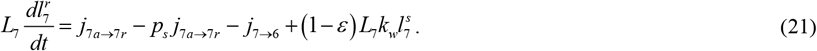

Eqs. (1)-(21) must be supplemented by the following equations for the mitochondria fluxes between the demand sites. *j*_*soma*→1_ is an input model parameter which characterizes the rate of mitochondria production in the soma (according to our model, they all enter the axon); equations for other anterograde fluxes (Fig. 2a) are

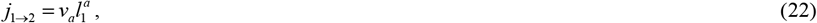

where *v*_*a*_ is the average velocity of motile mitochondria moving anterogradely.

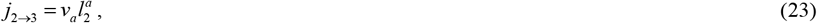

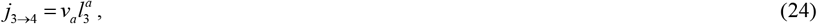

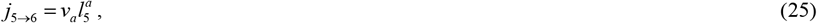

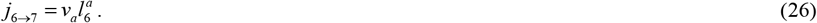

Equations for retrograde fluxes (Fig. 2a) are

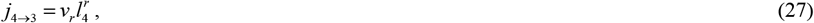

where *v*_*r*_ is the average velocity of motile mitochondria moving retrogradely.

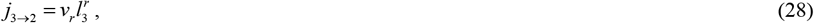

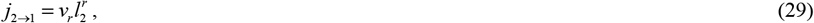

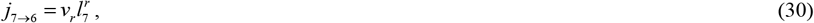

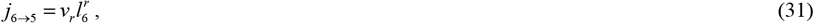

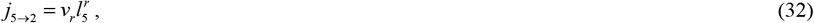

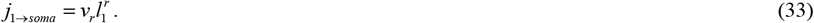

The turn-around fluxes (Fig. 2a) are simulated as follows:

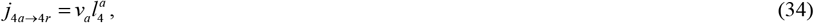

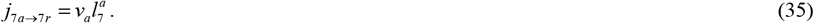

### 2.2. Model of mean age of mitochondria in the demand sites

To compute the age distributions of mitochondria in the stationary and moving states in the compartments surrounding the demand sites, the method described in [18] and [19] was utilized. In order to use the methodology described in the above papers, Eqs. (1)-(21) were restated as the following matrix equation:

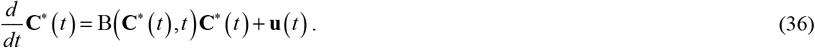

In Eq. (36), **C**^*^, using the terminology of [18], is the state vector, which is defined by its components:

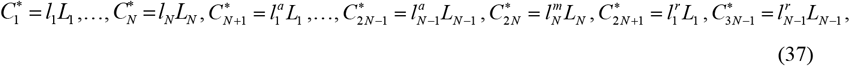

where

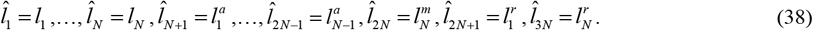

For the axons displayed in Figs. 1a, 1b, and 1c *N* =7, *N*=6, and *N* = 4, respectively. Although governing equations in the form given by (1)-(21) are easier to interpret in terms of mitochondria concentration in a compartment, rescaling these equations in the form given by Eqs. (36) and (37) is necessary because the conserved quantity is the total mitochondrial length in a certain compartment rather than its linear density (concentration). Also, restating the governing equations in the form given by Eqs. (36) and (37) enables the application of methods developed in [18] and [19] for computing the mean age and age density distributions of mitochondria in various demand sites.

The total length of mitochondria in stationary states is represented by the first *N* components of the state vector, the total length of mitochondria in anterograde states is represented by the next *N* components (*N*+1,…,2*N*), and the total length of mitochondria in retrograde states is represented by the last *N* components (2*N*+1,…,3*N*), see Fig. 2. Vector **u**(*t*) is defined below by Eqs. (102) and (103).

For the case of a branched axon with two unequal branches displayed in Fig. 1a, the elements of matrix B(3*N*,3*N*) (*N*=7) are given by Eqs. (39)-(101) below. We accounted for the internal mitochondria fluxes between the compartments, the external mitochondria flux entering the terminal from the axon, and the mitochondria flux leaving the terminal to move back to the axon (Fig. 2a). Equations for the following elements of matrix B defined in Eq. (36) were obtained by analyzing mitochondria fluxes to and from the compartments surrounding the most proximal demand site.

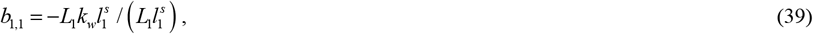

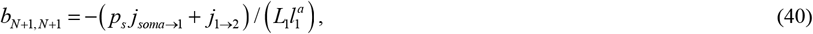

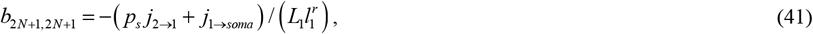

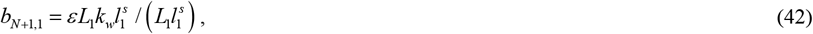

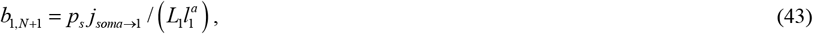

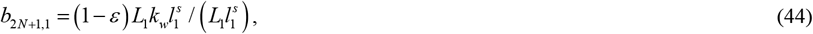

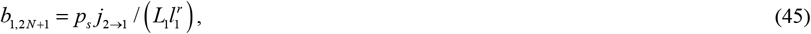

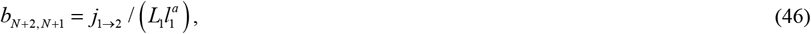

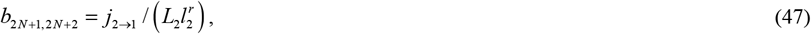

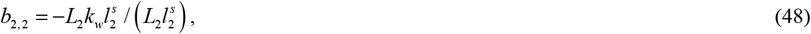

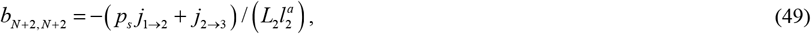

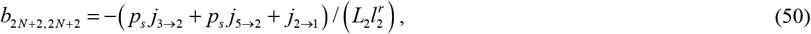

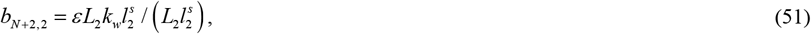

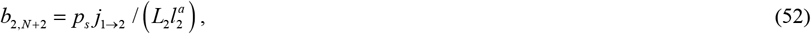

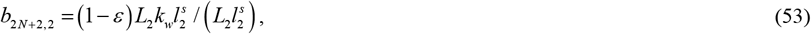

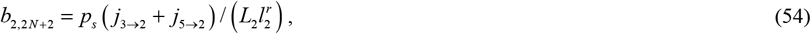

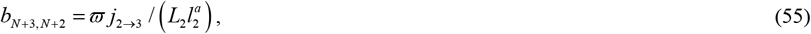

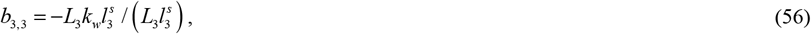

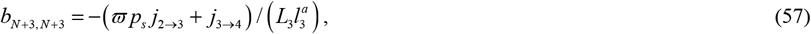

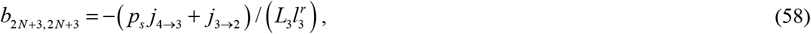

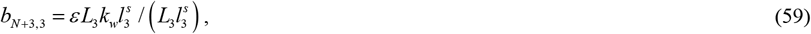

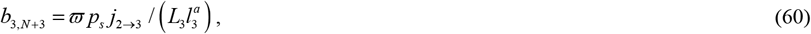

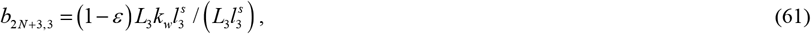

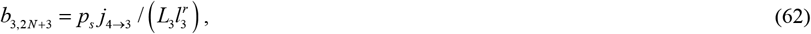

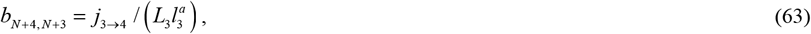

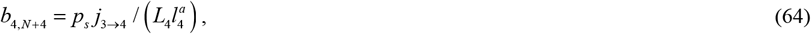

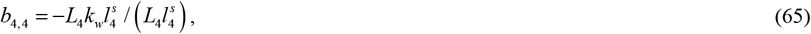

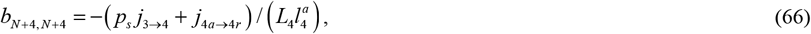

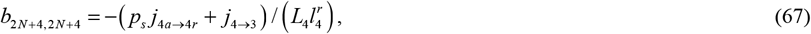

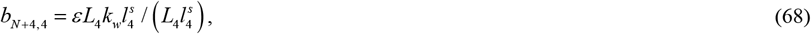

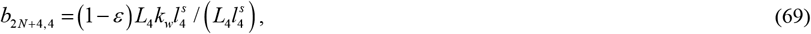

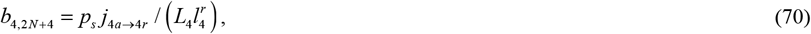

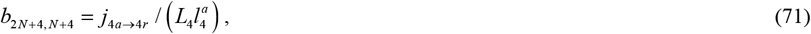

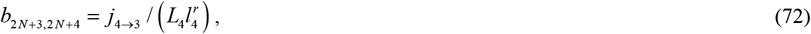

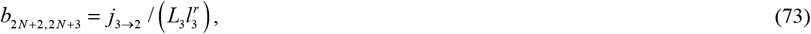

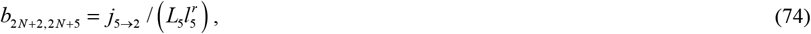

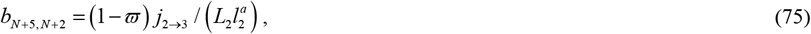

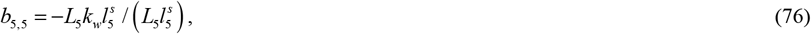

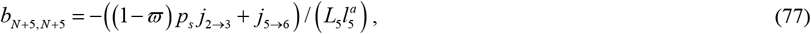

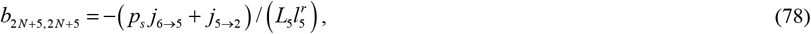

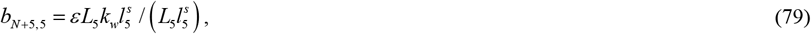

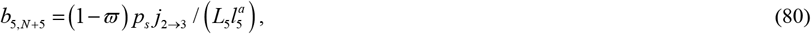

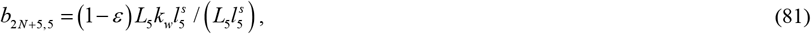

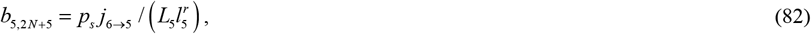

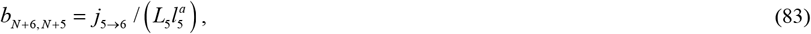

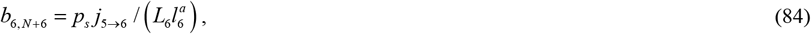

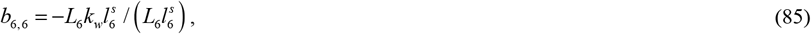

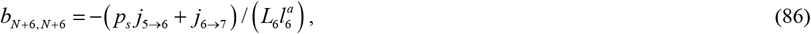

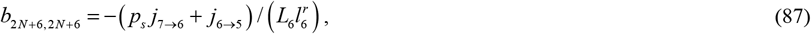

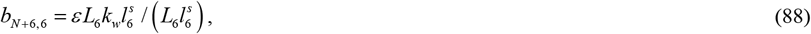

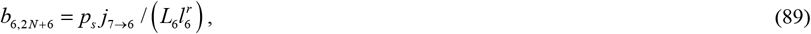

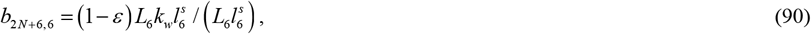

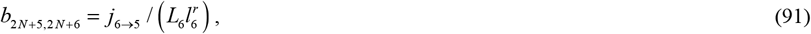

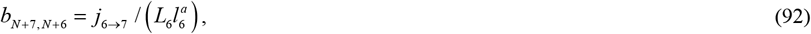

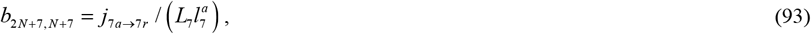

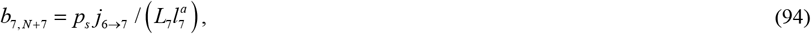

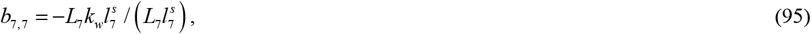

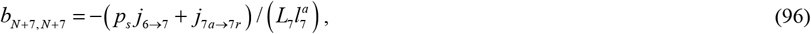

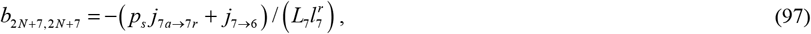

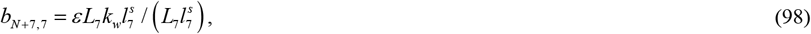

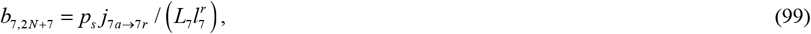

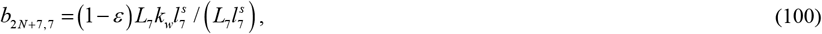

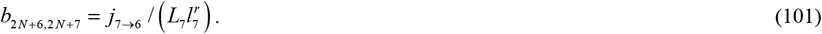

All other elements of matrix B, except for those given by Eqs. (39)-(101), are equal to zero. If a single mitochondrion in the axon is considered (its behavior is independent of all other mitochondria), the elements on the main diagonal control how long the mitochondrion stays in a certain compartment while off-diagonal elements determine the probabilities of its transition to other compartments [19].

The anterograde flux from the soma to the compartment with anterograde mitochondria by the most proximal demand site, *j*_*soma* →1_, is the only flux entering the axon. Our model assumes that all mitochondria that leave the axon (their flux is *j*_1→*soma*_) return to the soma for degradation, and none reenter the axon. We also assumed that mitochondria that enter the axon (their flux is *j*_*soma* →1_) are all newly synthesized in the soma and their age at the time of entry is equal to zero. The mitochondrial age calculated here should be understood as the time since mitochondria entered the axon.

The following equation yields the (*N*+1)^th^ element of vector **u**:

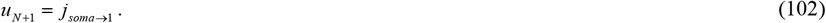

Because no other external fluxes enter the terminal,

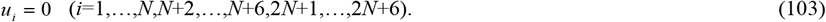

The state transition matrix Ф (effectively, a matrix-valued Green’s function), which is defined in [18], may be determined by solving the following matrix equation:

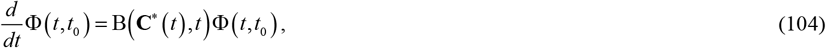

where *t*_0_is the initial time. In our case, *t*_0_ = 0.

The initial condition for Eq. (104) is

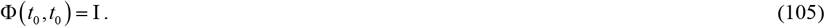

In Eq. (105), symbol I denotes an identity matrix.

We assumed that all mitochondria that enter the terminal are new. The age density of mitochondria entering the terminal after *t* = 0 can then be determined from the following equation:

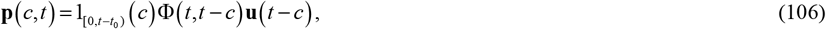

Where 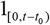 is the indicator function, which equals 1 if 0 ≤ *c* < *t* − *t*_0_ and equals 0 otherwise. The age density of mitochondria is the ratio of the total length of mitochondria in the compartment which have an age between *T* and *T+dT* over the duration of the interval *dT*. An integral of the age density of mitochondria over any time period with respect to time gives the length of mitochondria having an age within that time range.

The mean age of mitochondria in demand sites is [23]:

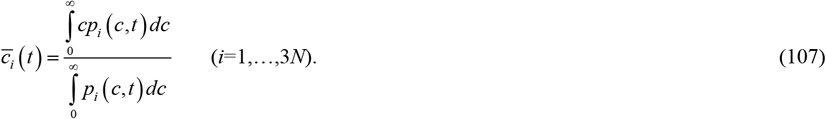

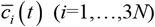 may be found by solving the following mean age system [18,23]:

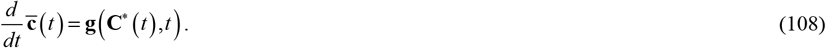

The initial condition for Eq. (108) is

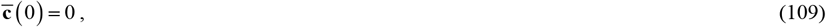

Where 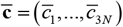. Vector **g** is defined as follows:

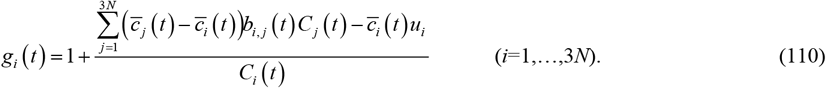

The following vectors were then defined to represent the mean age of mitochondria in the stationary, anterograde, and retrograde states in the demand sites, respectively:

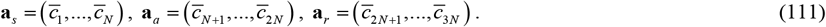

The arrays **a** _*s*_, **a** _*a*_, and **a**_*r*_ are one-dimensional and of size *N*.

Numerical solution of Eq. (108) was obtained utilizing MATLAB’s solver, ODE45. More details about the numerical solution are given in section S2 of Supplemental Materials. The results were also validated by using the formulas reported in [19]. In particular, the solution of Eq. (36) for a steady state can be obtained as:

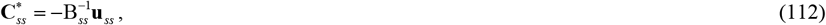

where the superscript −1 on a matrix denotes the inverse of the matrix and the subscript *ss* denotes steady state. The mean ages of mitochondria in various compartments displayed in Fig. 2 at steady state can be obtained as:

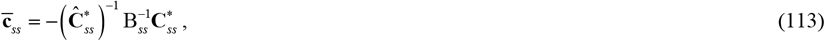

where

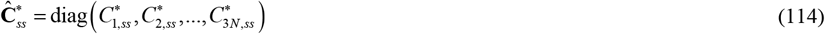

is the diagonal matrix with the components of the state vector, defined in Eq. (37), on the main diagonal. These components (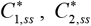, etc.) are calculated at steady state. Also in Eq. (113)

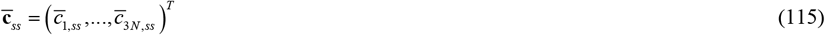

is the column vector composed of the mean ages of mitochondria in various compartments (superscript *T* means transposed).

The vector whose components represent age density of mitochondria residing in various compartments at steady state can be found as:

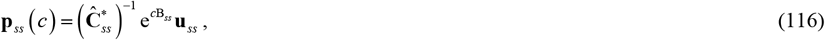

where e denotes the matrix exponential.

### 2.3. Sensitivity analysis

We have chosen the mean age of stationary mitochondria at steady state in the most distant demand site (#7, see Fig. 1a), *a*_*s*,7_, as a representative model output parameter. We investigated how *a*_*s*,7_ depends on model input parameters. This was done by computing the local sensitivity coefficients, which are first-order partial derivatives of *a*_*s*,7_ with respect to the model parameters [24-27]. For example, the sensitivity of *a*_*s*,7_ to parameter *p*_*s*_ was calculated as follows:

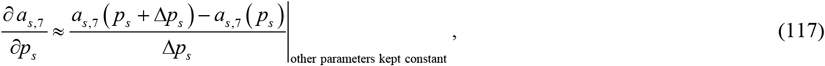

where Δ*p*_*s*_ = 10^−3^ *p*_*s*_ is the step size. We tested the independence of the sensitivity coefficients to the step size by varying the step sizes.

To make sensitivity independent of parameter magnitude, we followed [25,28] and calculated non-dimensional relative sensitivity coefficients as, for example:

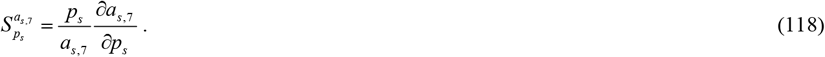

## 3. Results

We neglected the effect of mitophagy, and hence, at steady state, there is no reduction of mitochondria fluxes, except at the branching junction. If the flux of mitochondria splits equally between the shorter and longer branches (Fig. 3a, this situation is modeled by *ϖ*= 0.5), concentrations of mitochondria drop in half once the axon splits into two branches (Fig. 3b). This is explained as follows. We assumed that mitochondria enter the axon at a constant rate, (Fig. 2a,b). Anterograde fluxes are assumed to be proportional to the concentration of *j*_*soma*→1_ mitochondria in anterograde motile states (Eq. (22)-(26)). Because the anterograde velocity is assumed to remain the same, to simulate the reduction of mitochondria fluxes in branches the concentrations of mitochondria must also decrease by one half in the branches. The drop in mitochondria concentrations in the retrograde motile and stationary states is explained similarly. The drop in mitochondria concentration is consistent with [16], who pointed out that the density of mitochondria splits evenly at each branching junction.

**Fig. 3.**
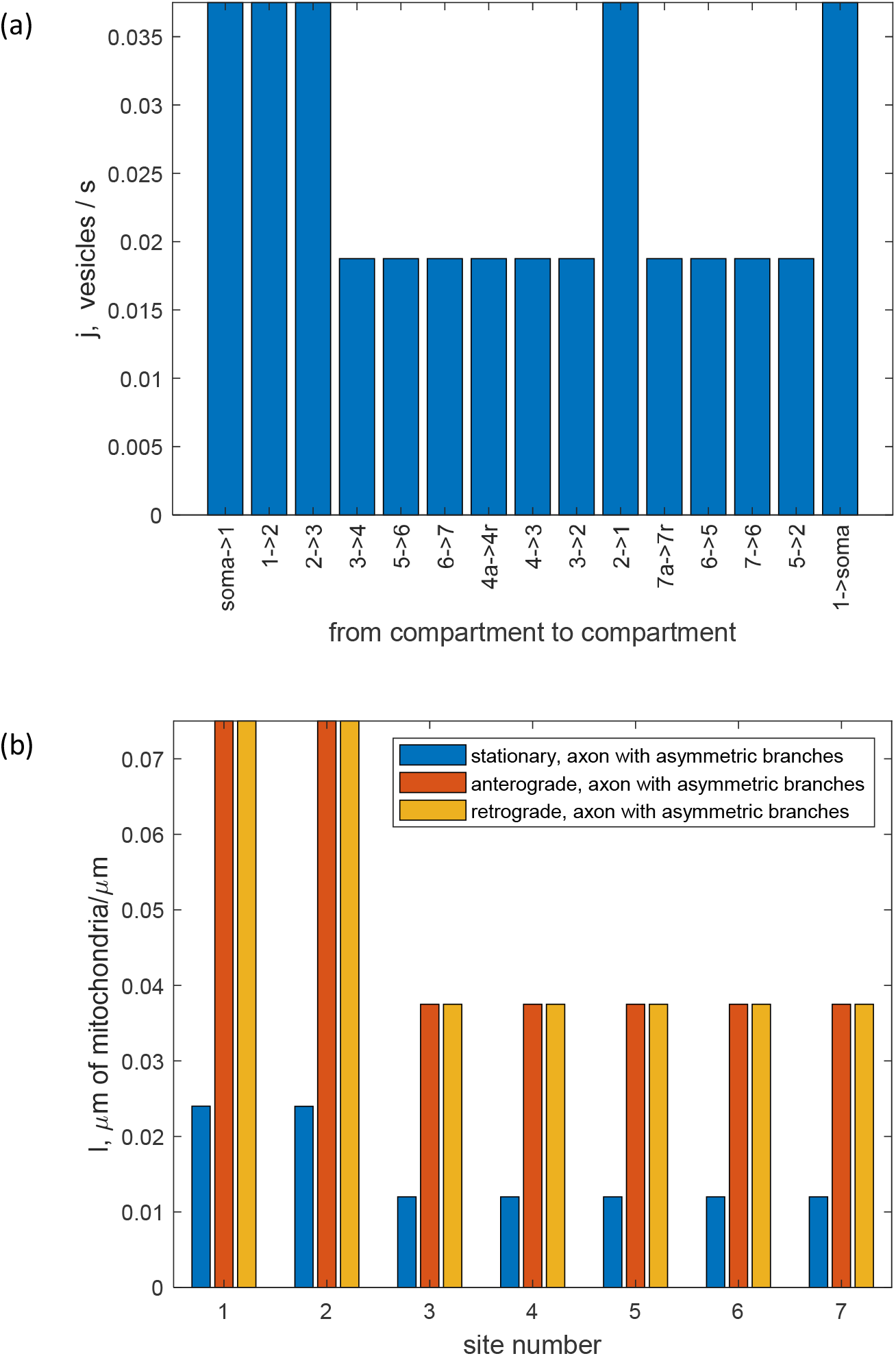
Branched axon with one shorter branch and one longer branch (Figs. 1a and 2a), steady state situation. (a) Fluxes of mitochondria traveling between the compartments. (b) Steady state values of the total length of stationary, anterogradely moving, and retrogradely moving mitochondria per unit length of the axon in the compartment by the *i*th demand site. *ϖ* = 0.5 (the flux of mitochondria splits equally between the shorter and longer branches).

The fluxes of mitochondria between the compartments for the situation when 1% of mitochondria at the branching junction enter the shorter branch and 99% enter the longer branch (*ϖ* = 0.01) are still constant in each branch (Fig. S1a) because no mitochondria destruction is assumed. The concentrations of mitochondria in a shorter branch that receives less mitochondria are much lower than in a longer branch (Fig. S1b).

In the axon with two unequal branches (Fig. 1a), the mean age of anterogradely moving mitochondria increases from the most proximal (#1) to the most distal (#4 or #7) demand site. This is as expected because mitochondria become older as they move from one demand site to the next demand site. Opposite to this, the age of retrograde mitochondria in the branches increases from the most distal demand site to the most proximal demand site with respect to the branching junction (#4 to #3 and #7 to #5). In the segment between the hillock and branching junction, the trend for the age of retrograde mitochondria is less clear because of the mixing of mitochondria coming from the two branches. The mean age of stationary mitochondria ranges between 10 and 16 hours. It generally increases from the hillock to the end of the branch, but with a lesser rate of increase than for anterogradely moving mitochondria. This is because mitochondria transition to the stationary state from the anterograde (containing younger mitochondria) and retrograde (containing older mitochondria) motile states (Fig. 4). The increase of the age of mitochondria residing in various demand sites with the distance from the soma is consistent with the experimental report in [29].

**Fig. 4.**
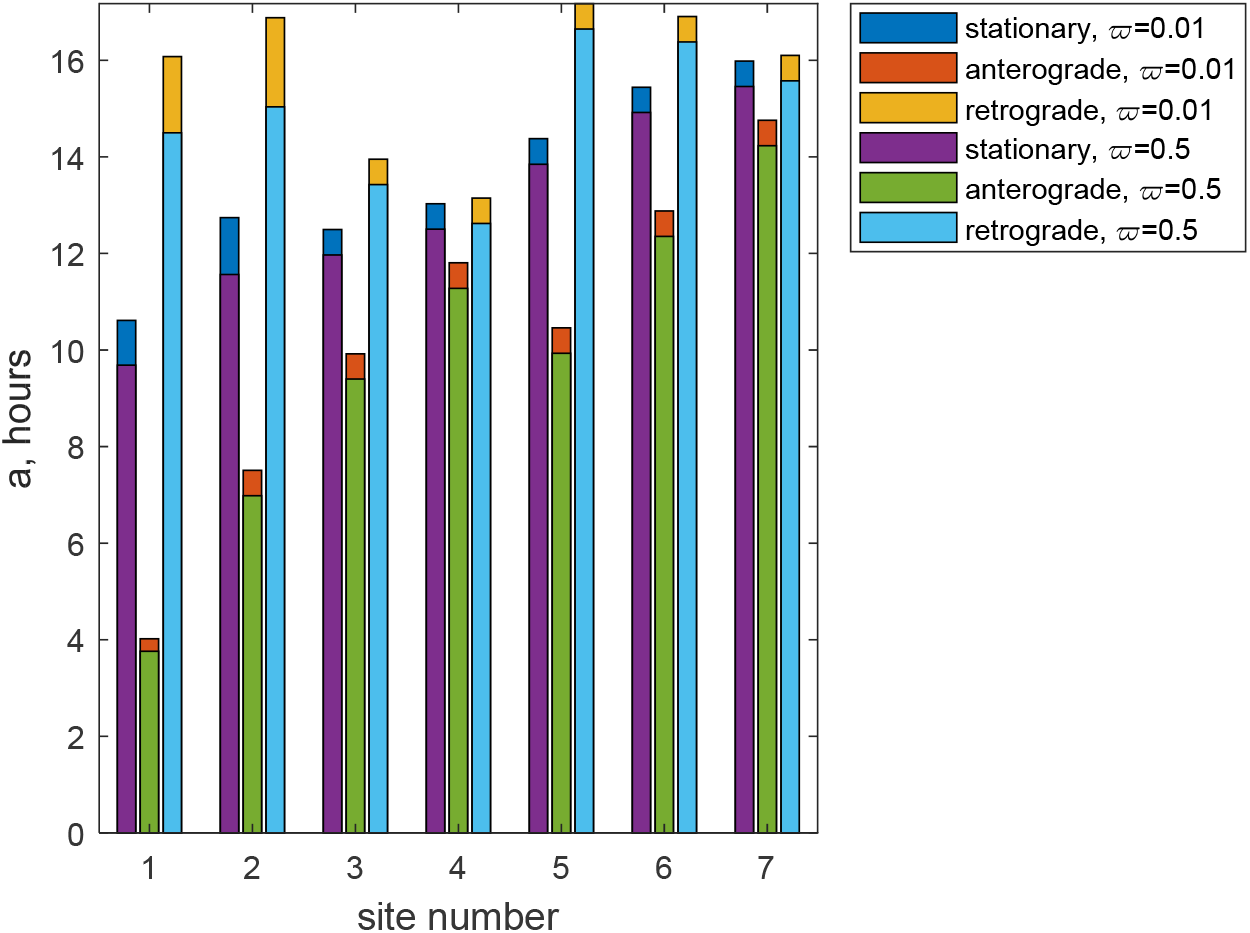
Branched axon with one shorter branch and one longer branch (Figs. 1a and 2a), steady state situation. Comparison of the mean age of stationary, anterogradely moving, and retrogradely moving mitochondria for the situation with *ϖ* = 0.01 (1% of mitochondria flux goes into the shorter branch and 99% goes into the longer branch) and *ϖ* = 0.5 (the mitochondria flux splits equally between the shorter and longer branches).

Noteworthy, the mean ages of mitochondria for an axon with two unequal branches do depend on how the flux of mitochondria splits at the branching junction. If 99% of anterogradely moving mitochondria go into the longer branch, *ϖ* = 0.01, the mean ages of mitochondria in all compartments are greater than if the mitochondria flux splits equally at the branching junction, *ϖ* = 0.5 (Fig. 4). This is because it takes mitochondria more time to travel to the end of the longer branch. Upon return to demand sites 2 and 1 (located before the branching junction), older mitochondria arriving from the longer branch mix with the rest of the mitochondria, thus elevating their mean age. Interestingly, this even elevates the mean age of mitochondria in the shorter branch (in demand sites 3 and 4). This is because older retrogradely moving mitochondria that return from the longer branch can transition to the stationary, and then to the anterograde states, then enter the shorter branch through the branching junction.

Ref. [29] used MitoTimer, a photoconvertible construct, which switches from green to red fluorescence over a period of hours due to slow oxidation as it ages. Using the results reported in [29], ref. [17] estimated the age of mitochondria at the axon tip to be 22 hours. The mean ages of mitochondria reported in Fig. 4 are consistent with this estimate. Our estimate is also consistent with [30] which reported that the complete turnover of axon terminal mitochondria in zebrafish neurons occurs within 24 h.

Fig. 5 displays the case when half of anterograde mitochondria enter the shorter branch, and half enter the longer branch at the branching junction (*ϖ* = 0.5). The range of the age densities of anterograde mitochondria is the narrowest, approximately between 0 and 20 hours (Fig. 5a). Newly synthesized anterograde mitochondria with zero age enter the anterograde compartment by the most proximal demand site (#1). This explains why the peak density of mitochondria occurs at zero age. The distribution of the age densities occurs because mitochondria spend some time pausing in this compartment, and also because there is an exchange between stationary and anterograde mitochondria (see the compartmental diagram in Fig. 2a). Stationary mitochondria, which are older, can again be brought to motion by kinesin motors. In terms of our model, this is interpreted as mitochondria reentering the anterograde motile state. If a stationary mitochondrion reenters the anterograde state, the average age of anterograde mitochondria increases. It is evident that in Fig. 5a the peaks for more distal demand sites are displaced in the direction of older mitochondria. This is because it takes time for mitochondria to travel to more distal demand sites.

**Fig. 5.**
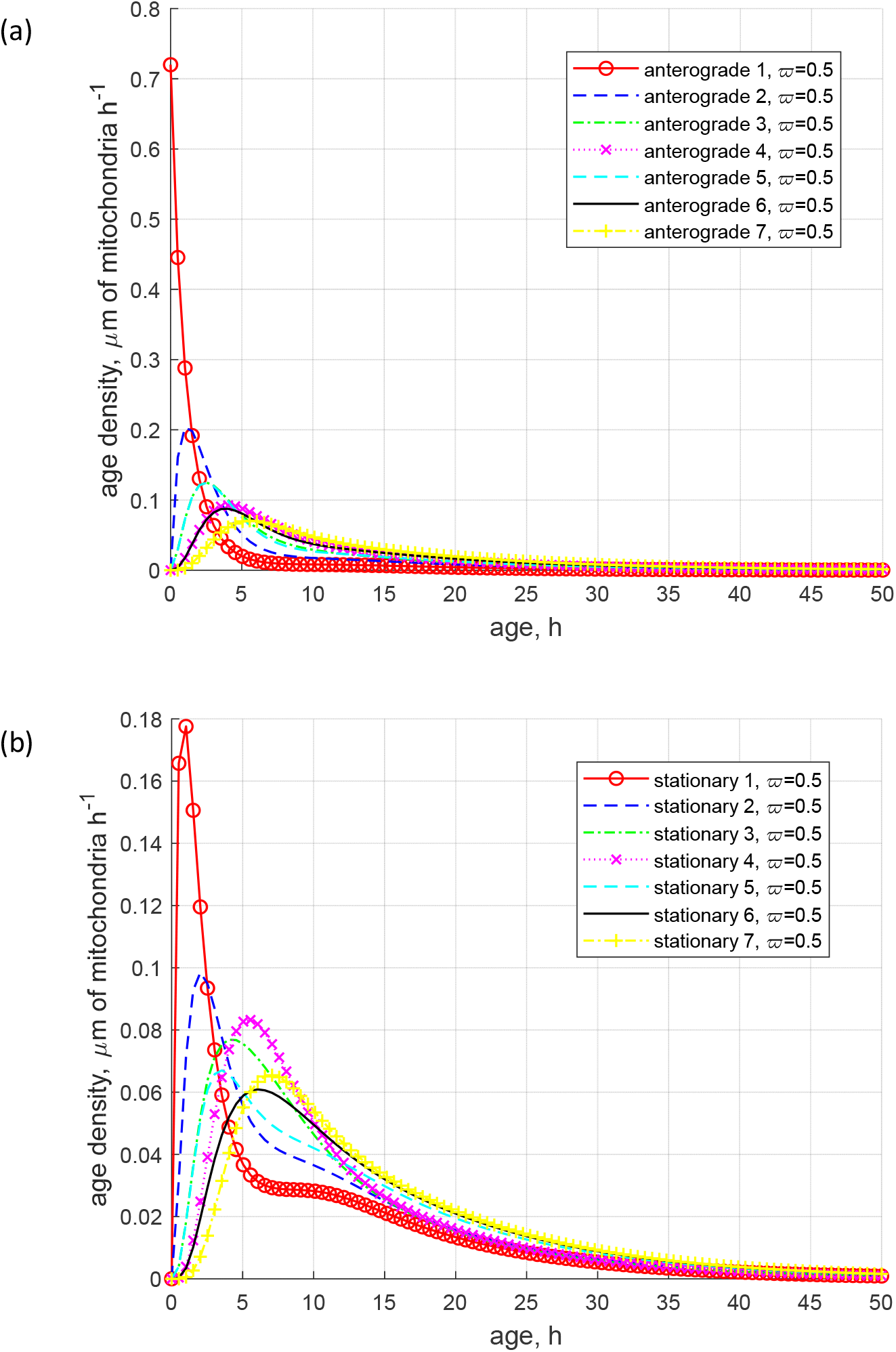
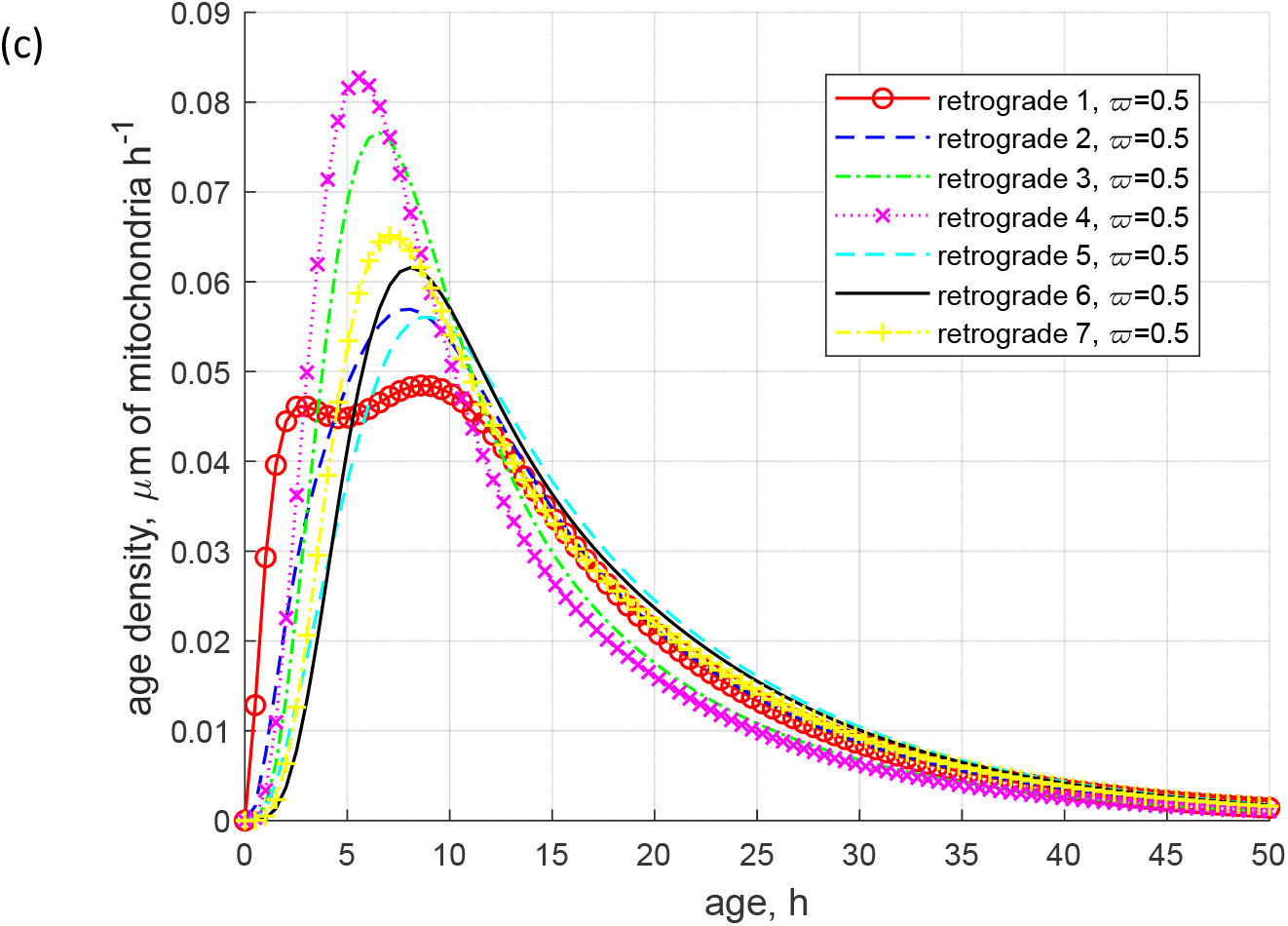
Branched axon with one shorter branch and one longer branch (Figs. 1a and 2a), steady state situation. (a) Age density of anterogradely moving mitochondria in various demand sites. (b) Age density of mitochondria in the stationary state in various demand sites. (c) Age density of retrogradely moving mitochondria in various demand sites. *ϖ* = 0.5 (the flux of mitochondria splits equally between the shorter and longer branches).

The range of the age density of stationary mitochondria is approximately between 0 to 40 hours (Fig. 5b). Thus, the age density distributions become more spread out than in Fig. 5a. This is explained by the fact that stationary states are fed by anterograde and retrograde states. Mitochondria in anterograde states are younger while mitochondria in retrograde states are older, and the mixing of younger and older mitochondria results in age density distributions that are more spread out. Similar to Fig. 5a, the peaks of the age density distributions are displaced toward older mitochondria for more distal demand sites.

The age density of retrograde mitochondria is even wider, approximately between 0 to 50 hours (Fig. 5c). The reason is that retrograde states contain old mitochondria that traveled all the way to the most distant demand sites and then turned around. Retrograde states also receive mitochondria transitioning from the stationary state, where the age density of mitochondria is already quite spread out. The locations of the peaks on the age density distributions are quite complicated. Opposite to what is observed for anterograde and stationary mitochondria, in general for retrograde mitochondria the peaks are displaced toward older mitochondria for more proximal demand sites. This is because retrograde mitochondria would generally turn around at the end of the branch, and they become even older as they travel from the end of the axon toward more proximal demand sites. In the retrograde compartment, at the most proximal demand site, a bimodal age distribution is observed. The peak corresponding to older mitochondria (which has a little higher magnitude in Fig. 5c) is attributed to mitochondria returning from their journey toward the ends of the branches. The peak corresponding to younger mitochondria (which has a little lower magnitude in Fig. 5c) is explained by mitochondria that just entered the most proximal demand site from the soma, transitioned to the stationary state, and then to the retrograde state.

As noted in [19], the presence of long tails in age density distributions means the presence of old particles in the system. The right-skewed age density distributions in Figs. 5a,b,c thus indicate that although the mean ages of mitochondria are of the order of 10 hours, there are also much older mitochondria in the axon, which stay in the terminal several times longer than that.

Fig. S2 displays the age densities of mitochondria for the case when 1% of anterograde mitochondria enter the shorter branch and 99% enter the longer branch at the branching junction (*ϖ* = 0.01). The difference in the age densities of anterograde mitochondria with the situation when mitochondria split equally at the branching junction (*ϖ* = 0.5) is minimal. The difference in the age densities of the stationary and retrograde mitochondria is only visible in the first two demand sites. The magnitudes of the peaks on the age density distribution of retrograde mitochondria in demand site 1 are now reversed: the peak corresponding to older mitochondria now has a lower magnitude while the peak corresponding to younger mitochondria has a higher magnitude (Fig. S2c). It is interesting that on the left-hand side of the curve corresponding to demand site 2, in the range of the mitochondria age between approximately 5 to 8 hours, the slope of the curve becomes less steep (Fig. S2c). While there is not a second peak at this location, the change in slope suggests that there is potential for one to form. This second peak would correspond to younger mitochondria. This reconfirms our explanation that younger mitochondria originate from anterograde mitochondria entering from the soma, transitioning to the stationary state (in this case corresponding to demand site 2), and then transitioning to the retrograde state.

To facilitate a comparison between the age density distributions for the case of *ϖ* = 0.01 and *ϖ* = 0.5, in the stationary states for the asymmetrically branched axon, we plotted age densities in the first demand sites in Fig. 6. It is interesting that for *ϖ* = 0.01 the curve displaying the age density distribution of stationary mitochondria in demand site 1 has a flat region between mitochondria ages of approximately 5 and 15 hours (Fig. 6a). The age density distributions corresponding to *ϖ* = 0.01 seem to have the centers of mass at lower ages, which would correspond to the lower mean ages of mitochondria. This, however, contradicts what is shown in Fig. 4. To explain this, in Fig. 5b we plotted the age densities in the range between 20 and 50 hours. Fig. 5b shows that the curves corresponding to *ϖ* = 0.01 are higher than the curves corresponding to *ϖ* = 0.5. This explains why the centers of mass of the age density distributions for *ϖ* = 0.01 are in realty displaced to the right of the centers of mass of the age density distributions for *ϖ* = 0.5 (all distributions in Fig. 6 are right skewed, but the distributions for *ϖ* = 0.01 have larger tails in the positive direction), which agrees with Fig. 4.

**Fig. 6.**
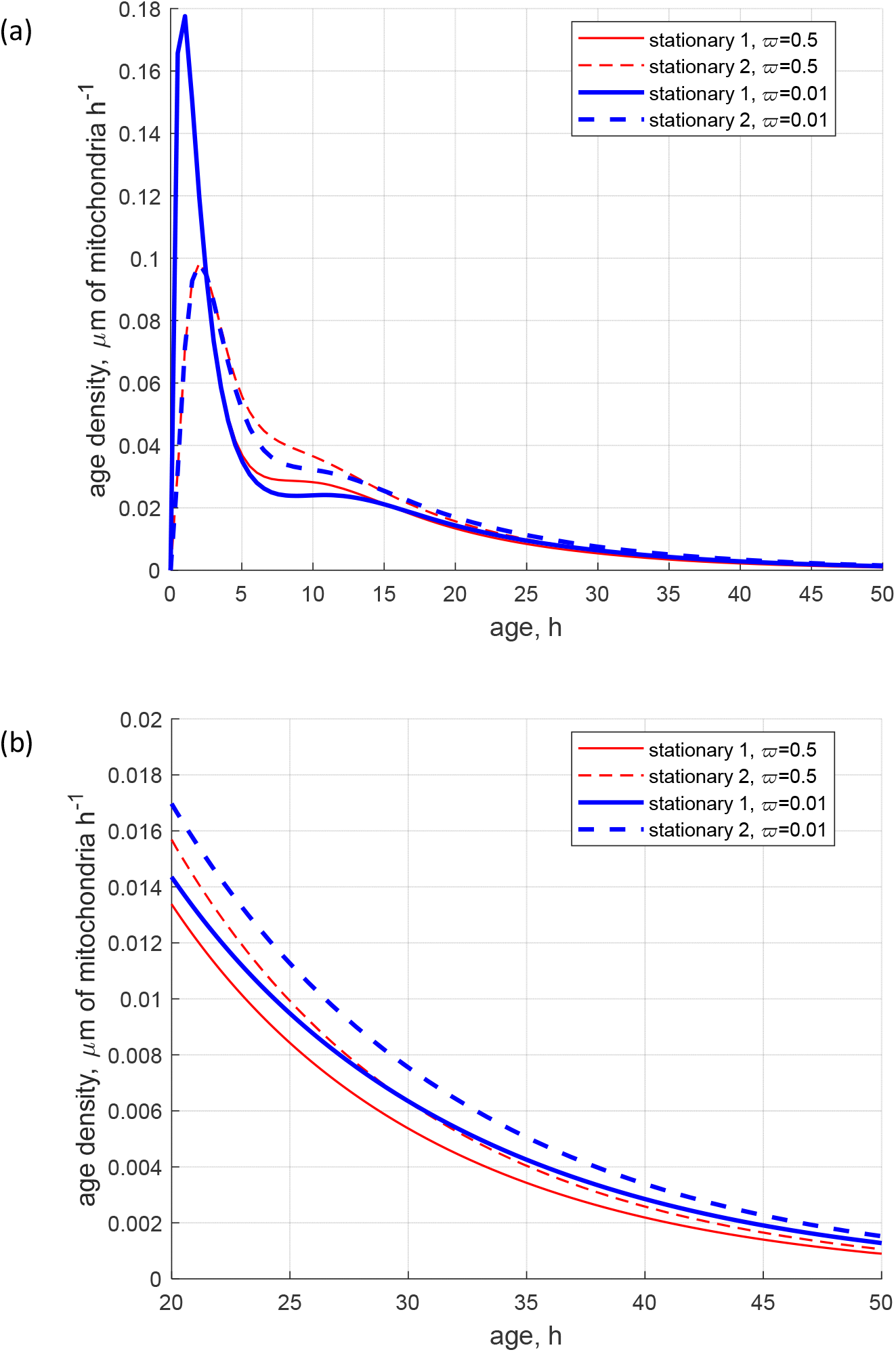
Branched axon with one shorter branch and one longer branch (Figs. 1a and 2a), steady state situation. Comparison between age density of mitochondria for the situation with *ϖ* = 0.5 (the mitochondria flux splits equally between the shorter and longer branches) and *ϖ* = 0.01 (1% of mitochondria flux goes into the shorter branch and 99% goes into the longer branch). (a) Age density of mitochondria in the stationary state in the first two (most proximal) demand sites. (b) Same as Fig. 6(a), but the range of the *x*-axis is between 20 and 50 hours.

It is interesting that changing the asymmetric axon to a symmetric one, with two identical branches, changes how the splitting of the flux of anterograde mitochondria at the branching junction affects the age of mitochondria. The axon with two identical branches and its compartmental representation are illustrated in Figs. 1b and 2b, respectively; it is obtained by removing the 7^th^ demand site from the longer branch in the asymmetric axon. Mitochondria fluxes between the compartments and mitochondria concentrations are shown in Figs. S3a and S3b, respectively. If the branches are identical, the way the mitochondria flux splits at the branching junction (characterized by *ϖ*) affects neither the mean ages of mitochondria (Fig. 7) nor their age density distributions (Fig. 8).

**Fig. 7.**
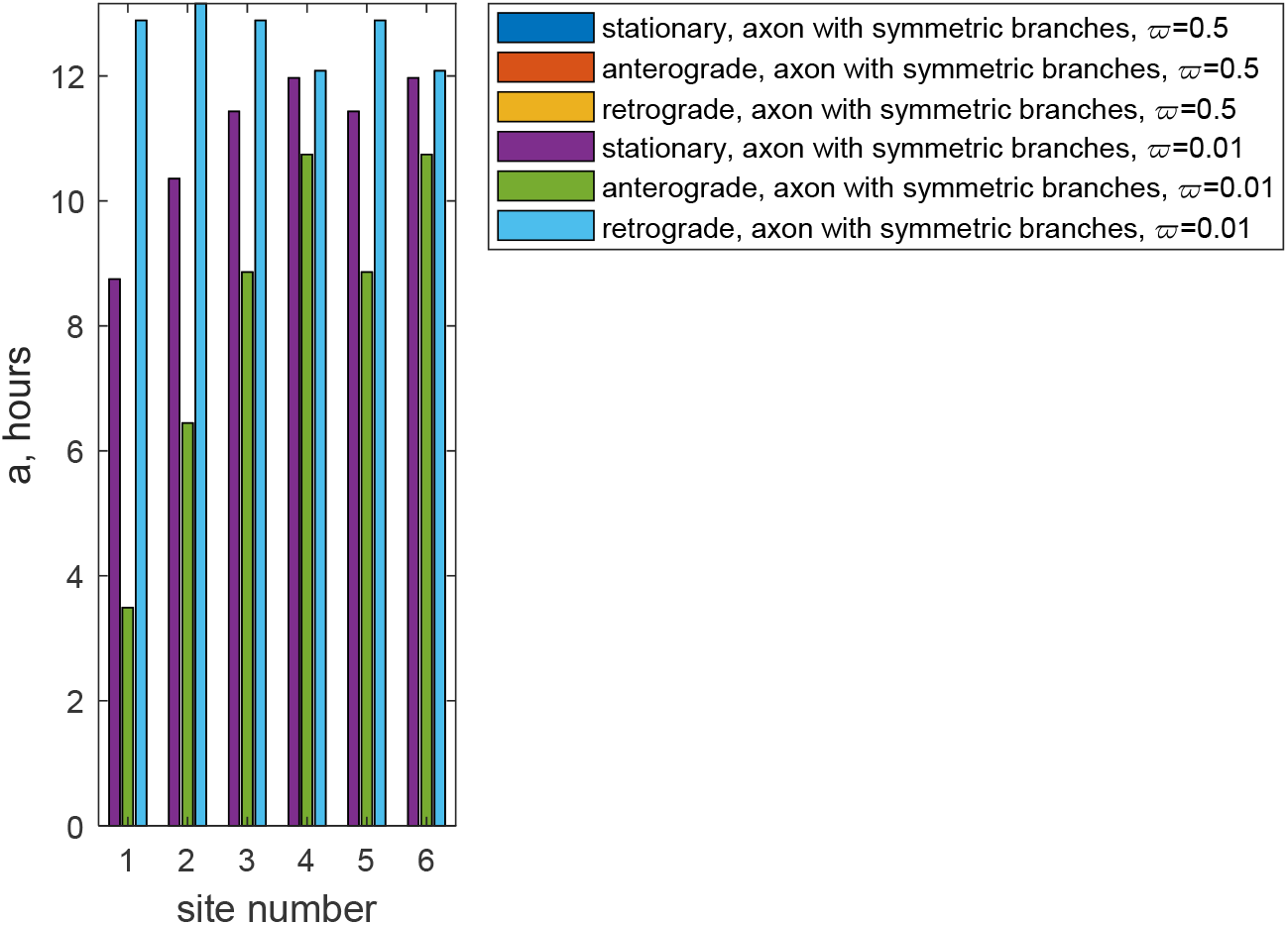
Branched axon with two equal branches (Figs. 1b and 2b), steady state situation. Comparison of the mean age of stationary, anterogradely moving, and retrogradely moving mitochondria for the situation with *ϖ* = 0.01 (1% of mitochondria flux goes into the upper branch and 99% goes into the lower branch) and *ϖ* = 0.5 (the mitochondria flux splits equally between the lower and upper branches).

**Fig. 8.**
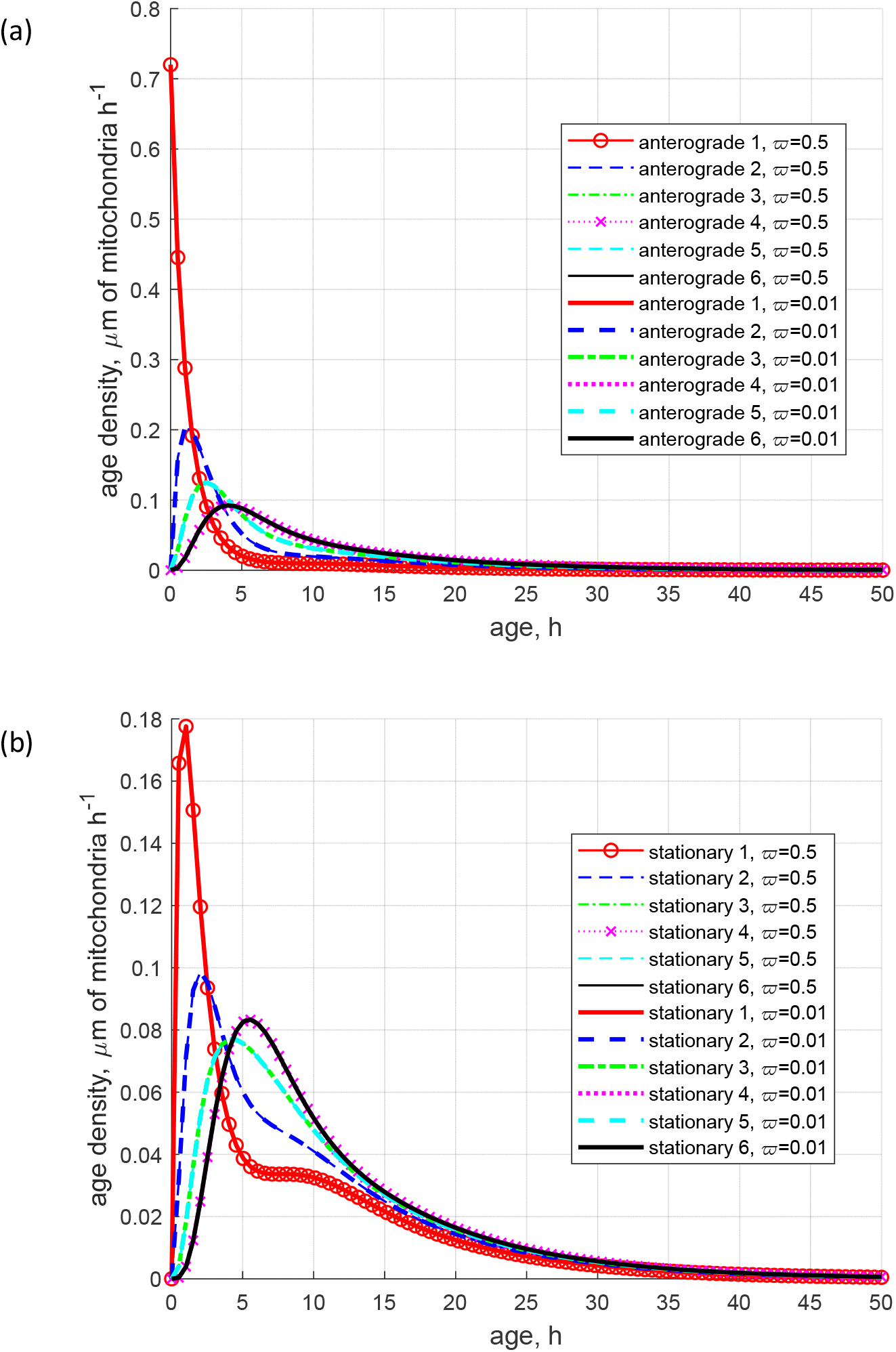
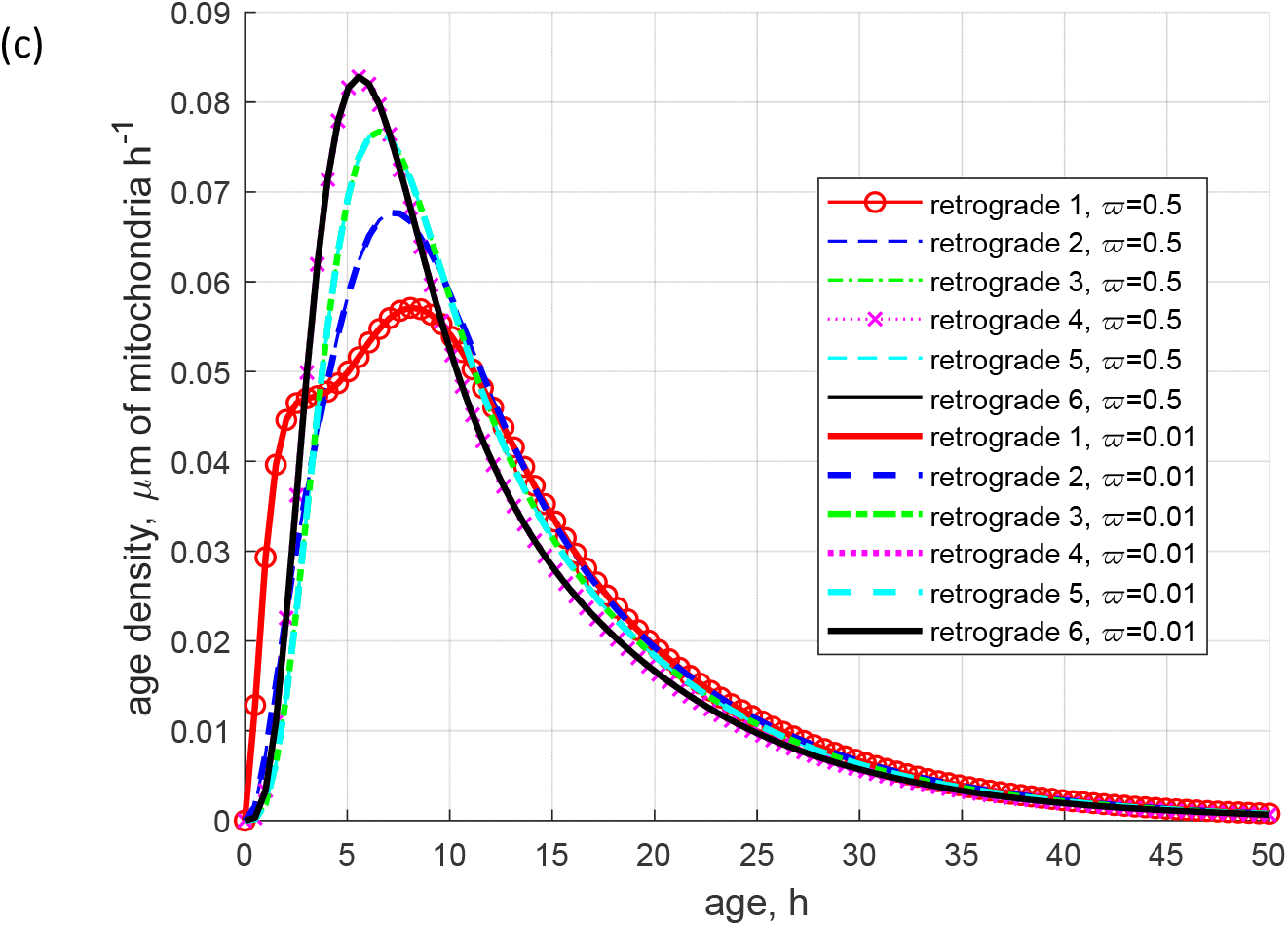
Branched axon with two identical branches (Figs. 1b and 2b), steady state situation. (a) Age density of anterogradely moving mitochondria in various demand sites. (b) Age density of mitochondria in the stationary state in various demand sites. (c) Age density of retrogradely moving mitochondria in various demand sites. Independence of the age density distributions of *ϖ* is shown by comparing the following two situations: *ϖ* = 0.5 (the fluxes of mitochondria are spit uniformly between the branches at the branching junction) and *ϖ* = 0.01 (1% of mitochondria flux goes into the shorter branch and 99% goes into the longer branch).

To further investigate this independence of *ϖ*, we studied the situation in a straight axon (Figs. 1c and 2c; the fluxes of mitochondria between the compartments in a straight axon are shown in Fig. S4a and mitochondria concentrations are shown in Fig. S4b). In terms of mitochondria transport, a straight axon displayed in Fig. 1c is identical to the symmetric axon displayed in Fig. 1b if *ϖ* = 1, that is all mitochondria enter one of the branches. A comparison in Fig. 9 shows that the mean ages of mitochondria in the four demand sites of the straight axon are identical to the mean ages of mitochondria in the first four demand sites of the symmetric axon (in the segment before the branching junction and in the upper branch). The age density distributions in these four demand sites of the straight and symmetric axons are also identical (data not shown).

**Fig. 9.**
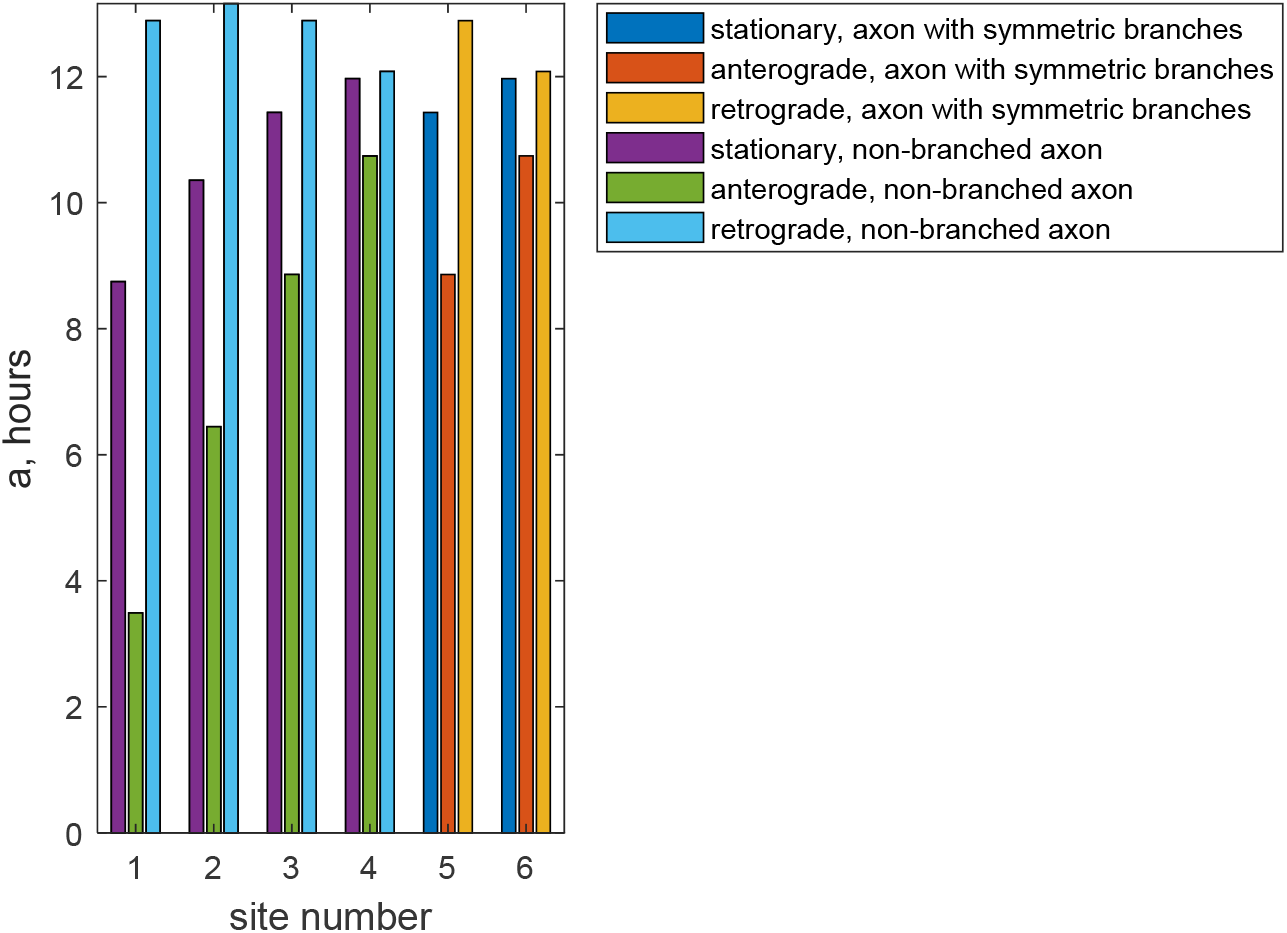
Comparison of the mean age of stationary, anterogradely moving, and retrogradely moving mitochondria in a branched axon with two symmetric branches (Figs. 1b and 2b) and in a non-branched (straight) axon. Computations show that the mean ages of mitochondria in all compartments in an axon with two symmetric branches are independent of the flux split ratio *ϖ*.

The fact that the mean age of mitochondria in demand sites # 1 and 2 is independent of whether the axon is branched is explained as follows. Imagine that mitochondria that enter the upper branch are colored red and that mitochondria that enter the lower branch are colored green. Because the problem is linear, nothing will change if all mitochondria (red and green) are directed into a single branch; their capture and release are independent of their color.

### 3.2. Investigating the sensitivity of the mean age of mitochondria to model parameters

We investigated the sensitivity of the mean stationary mitochondria age in demand site 7 (the most distal demand site), *a*_*s*,7_, to input model parameters. In Table 1, we report the relative sensitivity coefficients for *a*_*s*,7_ at steady state.

**Table 1.**
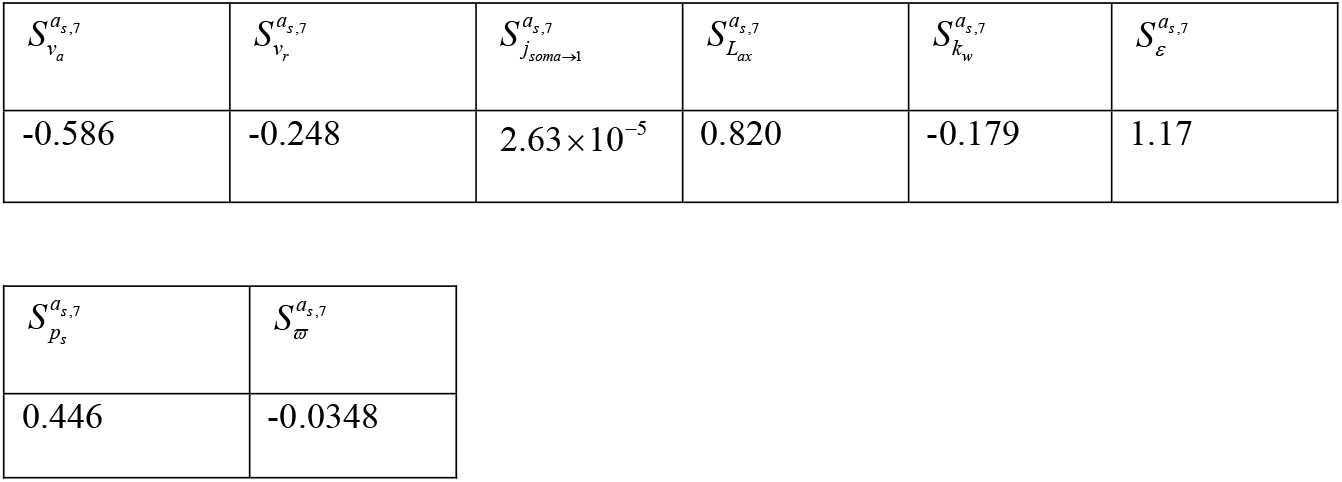
Relative sensitivity (defined in Eq. (118)) of the mean age of stationary mitochondria in demand site 7, *a*_*s*,7_, to various model parameters. Computations were performed using Eqs. (117) and (118) with (for example) Δ*p*_*s*_ = 10^−3^ *p*_*s*_ (very close results were obtained with Δ*p*_*s*_ = 10^−4^ *p*_*s*_ and Δ*p*_*s*_ = 10^−2^ *p*_*s*_). Computations were performed around the point with the following parameter values: *v*_*a*_ = 0.5 μms^-1^, *v*_*r*_ = 0.5 μms^-1^, *j*_*soma*→1_ = 0.0375 mitochondria s^-1^, *L*_*ax*_ = 10^4^ μm, *k*_*w*_ = 5 ×10^−4^ s^-1^, *ε* = 0.5, *p*_*s*_ = 0.4, and *ϖ* = 0.5.

Out of eight model parameters, *a*_*s*,7_ is most sensitive to parameter *ε*. This parameter characterizes how mitochondria are split between the anterograde and retrograde pools when they are released from the stationary pool. The relative sensitivity 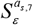 is positive, which means that *a*_*s*,7_ increases when *ε* is increased. This is because mitochondria released into the anterograde pool spend more time in the terminal and have a higher chance of capture and inclusion in the stationary pool. *a*_*s*,7_ also exhibits positive sensitivity to the length of the axon, *L*_*ax*_, and to the probability for a motile mitochondrion to be captured into the stationary state, *p*_*s*_. The increase of *p*_*s*_ leads to mitochondria spending more time in the stationary pool. Interestingly, *a*_*s*,7_ is almost independent of the rate of mitochondria production in the soma, *j*_*soma*→1_. *a*_*s*,7_ exhibits the largest negative sensitivity to the average velocity of anterograde motion of mitochondria, *v*_*a*_, which means that an increase of *v*_*a*_ decreases *a*_*s*,7_. This is expected because faster transport velocity decreases the average time mitochondria spend in the terminal. Similarly, *a*_*s*,7_ exhibits negative sensitivity to the average velocity of retrograde motion of mitochondria, *v*_*r*_. The sensitivity of *a*_*s*,7_ to the kinetic constant that characterizes the rate of stationary mitochondria reentry into the motile states, *k*_*w*_, is also negative. This is also expected because for smaller values of *k*_*w*_ mitochondria stay longer in the stationary state. The sensitivity of *a*_*s*,7_ to the parameter that indicates what portion of mitochondria enter the shorter branch, *ϖ*, is also slightly negative.

## 4. Discussion, limitations of the model, and future directions

We investigated the effects of axon splitting into two asymmetric or symmetric branches on mitochondria fluxes and concentrations, mitochondria mean age, and age density distribution. When the axon splits, the flux of anterogradely running mitochondria divides between the branches. Because fewer mitochondria move in the branch than in the segment of the axon before the branching junction, the concentration of mitochondria in the branch is less than in the segment of the axon before the branching junction. For example, if the flux of mitochondria divides equally at the branching junction, the concentrations of mitochondria in the branches reduce by a factor of two in the branches. Since DA axons branch multiple times, this shows that the mitochondria concentrations in distal branches can be very low.

However, if the axon splits into two equal branches, the mean age of mitochondria and the age density distributions are not affected by axon splitting. Furthermore, these quantities are independent of what portion of mitochondria go into one branch and what go into the other branch. Even if all mitochondria at the branching junction enter one branch (which in terms of mitochondria transport is equivalent to a non-branching (straight) axon), the mean age and age density of mitochondria remain the same. The independence of mitochondria age (which should be directly correlated with mitochondrial health) on branching may explain how DA neurons are able to support their enormous arbors. Although many mitochondria proteins can be replaced locally, either by mitochondria fusion or by local synthesis of mitochondria proteins [31], older mitochondria generally have lower membrane potential, and therefore are less healthy, and have to be transported to the soma for recycling in lysosomes or for servicing via fusion with newly synthesized mitochondria [30,32].

Our simulations predict that the age density of retrograde motile mitochondria in the proximal region of the axon close to the axon hillock has a distribution characterized by two peaks (Fig. 5c). The peak corresponding to older mitochondria is attributed to mitochondria returning after a long trip to the end of axon branches. The peak corresponding to younger mitochondria is attributed to new mitochondria that were just synthesized in the soma, transitioned to the stationary state, and then were re-released into the retrograde state. The study of the age of retrogradely moving mitochondria in the proximal axonal region (which can be done by using a photoconvertible construct, such as MitoTimer [29]) can provide information about the mechanism of mitochondria re-release from the stationary into the motile states. If mitochondria are re-released randomly, with a 50-50% chance of going either to the anterograde or to the retrograde state (as our simulations assume; the same assumption was also used in [16]), the age density of retrograde mitochondria close to the hillock should have a distribution with two peaks, as shown in Fig. 5c. If, however, such distribution is not observed, this indicates that the choice of whether mitochondria are re-released into the anterograde or retrograde pool is made based on the age of mitochondria: young mitochondria are more likely to be re-released into the anterograde pool and old mitochondria are more likely to be re-released into the retrograde pool.

In summary, the answers to the four questions of paragraph 5 in the Introduction are

1. When an axon splits into two branches, mitochondria concentrations after the branching junction decrease. The reduction depends on how the flux of mitochondria splits at the branching junction. For example, if the flux of mitochondria splits equally between the branches, the concentrations of mitochondria drop in half.
2. When an axon splits into two equal branches, the mean age of mitochondria is not affected. The ages of mitochondria in the demand sites are the same as in a non-branching axon of the same length.
3. If the axon splits into two unequal branches, the situation becomes more complicated because of the mixing of older retrograde mitochondria returning from the longer branch and younger retrograde mitochondria returning from the shorter branch. The mixing of older and younger mitochondria affects age density distributions in the demand sites before and after the branching junction.
4. If the axon splits into two equal branches, the mean ages and age densities of mitochondria do not depend on how the mitochondria flux splits at the branching junction. If the axon splits into two unequal branches, it does matter what portion of mitochondria enter the shorter branch and what portion enter the longer branch. This is because in more distal demand sites mitochondria are older, and if a larger portion of mitochondria enter the longer branch, mitochondria in other demand sites will also be older due to the mixing with retrogradely transported mitochondria.

A connection exists between changes in axon morphology and nervous system disorders [33]. Future development of the model should involve investigating transient situations, which may result, for example, as suggested in [16], from changing metabolic needs. This can be simulated, for example, by changing the value of parameter *ϖ*, which simulates how the mitochondria flux splits between the two branches.

Our current model assumes that all mitochondria entering the axon from soma are new (have zero age). However, it is likely that retrograde mitochondria returning to the soma are not destroyed in lysosomes, but are serviced by fusion with newly synthesized mitochondria, which would renew their proteins [30]. The renewed mitochondria are then returned to the soma. Future versions of the model should account for this possibility. Neglecting mitochondria clearance by mitophagy is also a limitation of the current model, which should be addressed in future research.

The model developed in this paper cannot distinguish between lengths of individual mitochondria. All mitochondria located in the same compartment are lumped together and are represented by a single parameter called the total length of mitochondria. Future models should overcome this limitation. This would require modeling fusion and fission of mitochondria.

In this paper, we have chosen the model parameters based on the computational study of [16], which optimized the parameters based on mitochondrial health. Investigating model behavior for a wider range of parameters, and the sensitivity of the solutions to more parameters, should be performed in future work.

## Acknowledgements

IAK acknowledges the fellowship support of the Paul and Daisy Soros Fellowship for New Americans and the NIH/National Institute of Mental Health (NIMH) Ruth L. Kirchstein NRSA (F30 MH122076-01). AVK acknowledges the support of the National Science Foundation (award CBET-2042834) and the Alexander von Humboldt Foundation through the Humboldt Research Award.

## Ethical Statement

None.

## Data availability statement

This article has no additional data.

## Authors’ contributions

IAK and AVK contributed equally to the performing of computational work and article preparation. Both authors approved the final version of the manuscript and agreed to be accountable for all aspects of the work.

## Supplemental Materials

### S1. Supplementary tables

**Table S1.** Dependent variables in the compartmental model of mitochondria transport in the axon.

| Symbol | Definition | Units |
| --- | --- | --- |
| $l_i^a$ | Concentration (per unit length of the axon) of anterograde motile mitochondria in the compartment by the $i$th demand site | ($\mu\text{m}$ of mitochondria)/$\mu\text{m}$ |
| $l_i^s$ | Concentration (per unit length of the axon) of stationary mitochondria in the compartment by the $i$th demand site | ($\mu\text{m}$ of mitochondria)/$\mu\text{m}$ |
| $l_i^r$ | Concentration (per unit length of the axon) of retrograde motile mitochondria in the compartment by the $i$th demand site | ($\mu\text{m}$ of mitochondria)/$\mu\text{m}$ |
| $j_{a \rightarrow b}$ | Flux of mitochondria from compartment $a$ to compartment $b$ which is represented by the total length of mitochondria transported from compartment $a$ to compartment $b$ per unit time | ($\mu\text{m}$ of mitochondria) $\text{s}^{-1}$ |

**Table S2.**
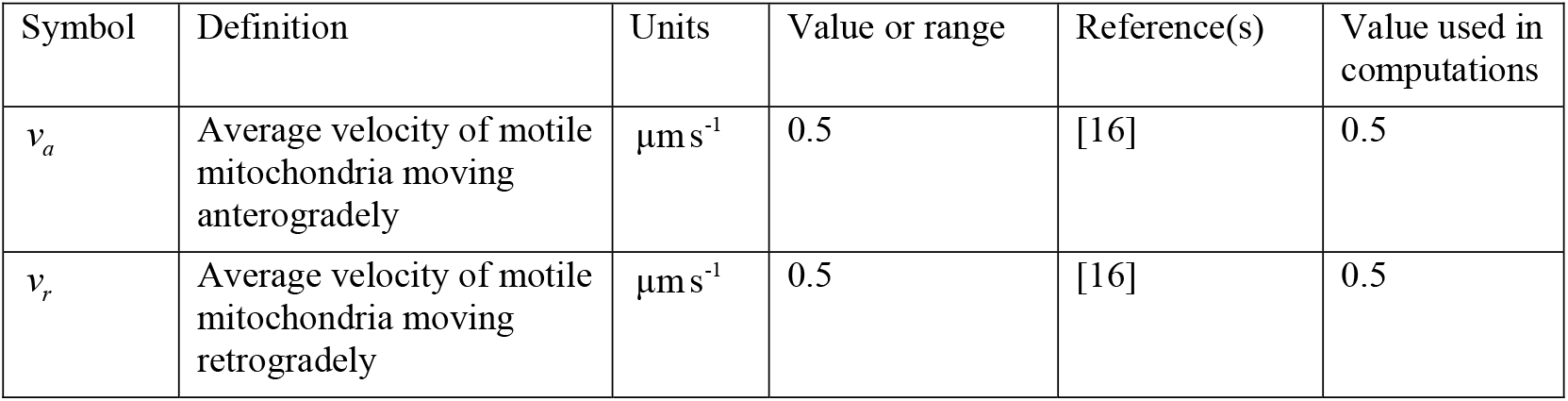
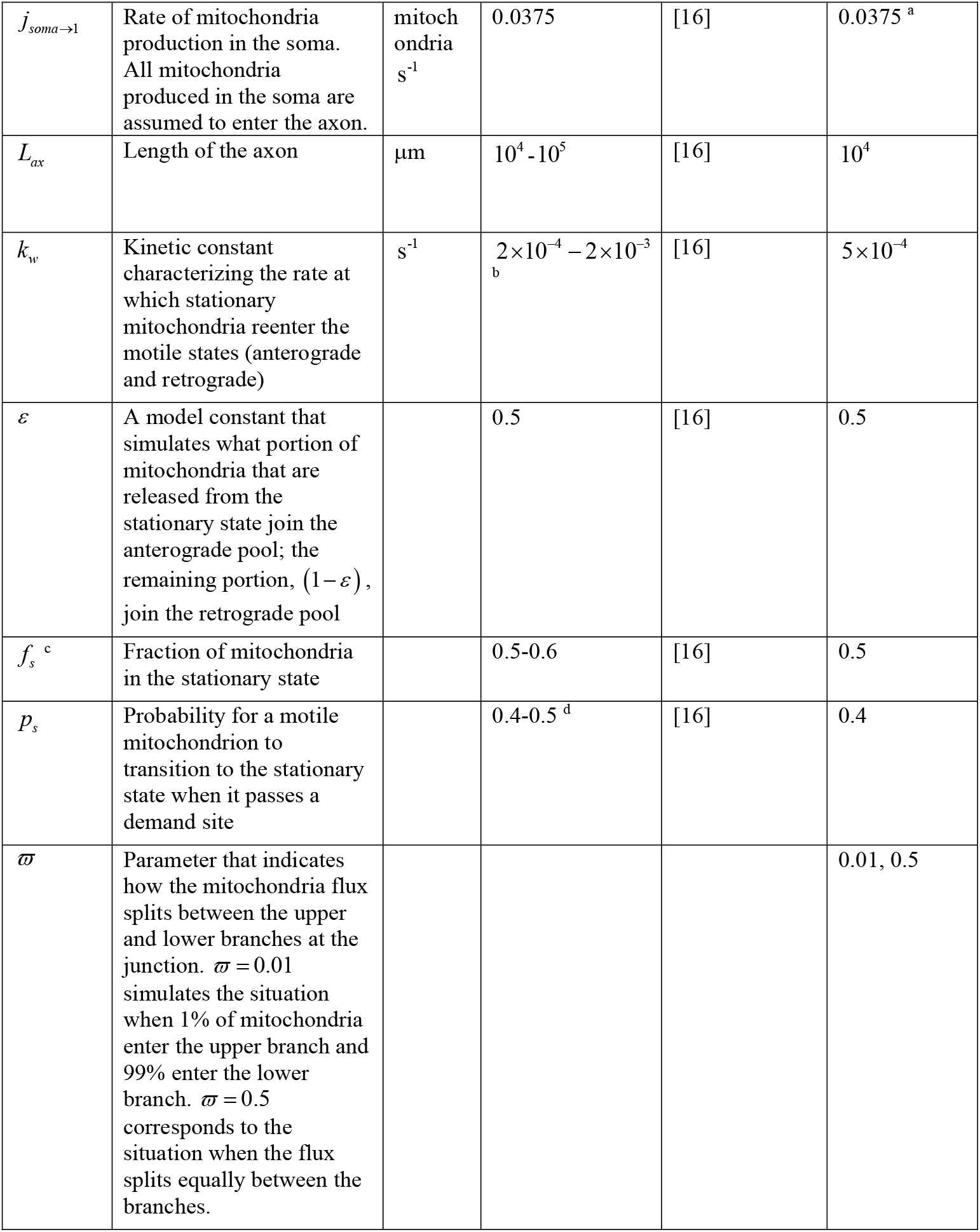
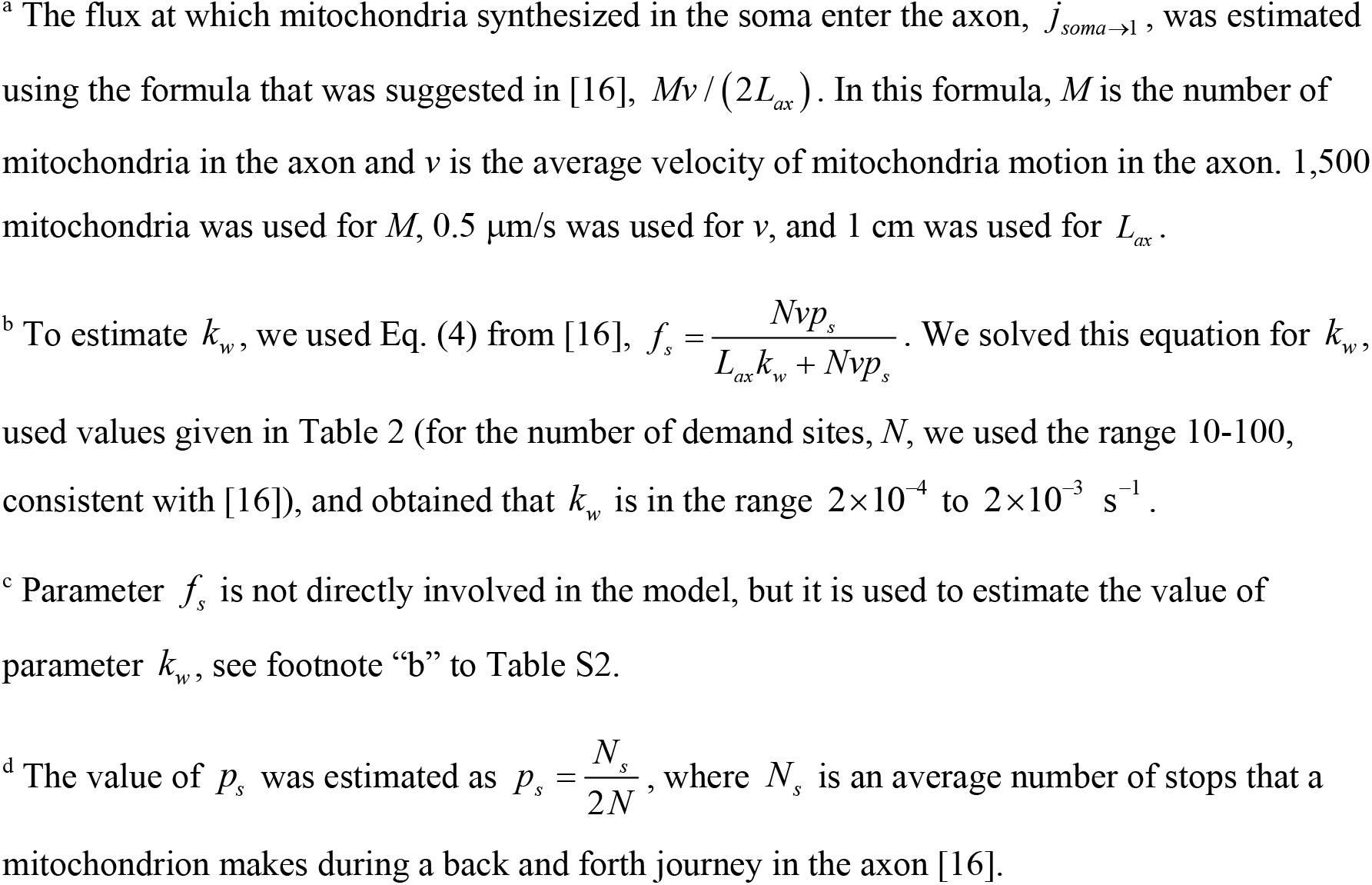
Parameters characterizing mitochondrial transport model in the axon.

| Symbol | Definition | Units | Value or range | Reference(s) | Value used in computations |
| --- | --- | --- | --- | --- | --- |
| $v_a$ | Average velocity of motile mitochondria moving anterogradely | $\mu\text{m s}^{-1}$ | 0.5 | [16] | 0.5 |
| $v_r$ | Average velocity of motile mitochondria moving retrogradely | $\mu\text{m s}^{-1}$ | 0.5 | [16] | 0.5 |
| $j_{soma \rightarrow 1}$ | Rate of mitochondria production in the soma. All mitochondria produced in the soma are assumed to enter the axon. | mitochondria s<sup>-1</sup> | 0.0375 | [16] | 0.0375<sup>a</sup> |
| $L_{ax}$ | Length of the axon | μm | 10<sup>4</sup>-10<sup>5</sup> | [16] | 10<sup>4</sup> |
| $k_w$ | Kinetic constant characterizing the rate at which stationary mitochondria reenter the motile states (anterograde and retrograde) | s<sup>-1</sup> | $2 \times 10^{-4} - 2 \times 10^{-3}$<sup>b</sup> | [16] | $5 \times 10^{-4}$ |
| $\varepsilon$ | A model constant that simulates what portion of mitochondria that are released from the stationary state join the anterograde pool; the remaining portion, $(1 - \varepsilon)$, join the retrograde pool | | 0.5 | [16] | 0.5 |
| $f_s$<sup>c</sup> | Fraction of mitochondria in the stationary state | | 0.5-0.6 | [16] | 0.5 |
| $p_s$ | Probability for a motile mitochondrion to transition to the stationary state when it passes a demand site | | 0.4-0.5<sup>d</sup> | [16] | 0.4 |
| $\varpi$ | Parameter that indicates how the mitochondria flux splits between the upper and lower branches at the junction. $\varpi = 0.01$ simulates the situation when 1% of mitochondria enter the upper branch and 99% enter the lower branch. $\varpi = 0.5$ corresponds to the situation when the flux splits equally between the branches. | | | | 0.01, 0.5 |

<sup>a</sup> The flux at which mitochondria synthesized in the soma enter the axon, $j_{soma \rightarrow 1}$ , was estimated using the formula that was suggested in [16], $Mv / (2L_{ax})$ . In this formula, $M$ is the number of mitochondria in the axon and $v$ is the average velocity of mitochondria motion in the axon. 1,500 mitochondria was used for $M$ , 0.5 $\mu\text{m/s}$ was used for $v$ , and 1 cm was used for $L_{ax}$ .
<sup>b</sup> To estimate $k_w$ , we used Eq. (4) from [16], $f_s = \frac{Nvp_s}{L_{ax}k_w + Nvp_s}$ . We solved this equation for $k_w$ , used values given in Table 2 (for the number of demand sites, $N$ , we used the range 10-100, consistent with [16]), and obtained that $k_w$ is in the range $2 \times 10^{-4}$ to $2 \times 10^{-3} \text{ s}^{-1}$ .
<sup>c</sup> Parameter $f_s$ is not directly involved in the model, but it is used to estimate the value of parameter $k_w$ , see footnote “b” to Table S2.
<sup>d</sup> The value of $p_s$ was estimated as $p_s = \frac{N_s}{2N}$ , where $N_s$ is an average number of stops that a mitochondrion makes during a back and forth journey in the axon [16].

### S2. Numerical solution procedure

In this paper, we presented two alternative methods for investigating the steady state mitochondria distributions in the axon, which we used to validate each other. The first method is based on solving transient equations until they reach steady state. Since we are only interested in the solution at steady state, we can choose any initial condition (for simplicity, we used zero boundary condition). Eqs. (1)-(21), simulating mitochondria transport in the axon with two unequal branches (Fig. 1a), were solved using MATLAB’s solver, ODE45 (MATLAB R2020b, MathWorks, Natick, MA, USA). The same was done for the axon with two equal branches depicted in Fig. 1b (Eqs. (1)-(15) and (S1)-(S3)) and for the straight axon depicted in Fig. 1c (Eqs. (1)-(3) and (S12)-(S20)). Eq. (108) for the mean ages of mitochondria with initial condition (109) was also solved by ODE45 until the mean ages reached their steady state values. Although in this paper the above equations were used to investigate the solution at steady state, the developed transient equations can be also used to investigate mitochondria redistribution during branch growth, modifying activity patterns, or axotomy [4].

Eqs. (112)-(116) allow finding steady state solutions directly. We implemented these equations using standard MATLAB operators, such as matrix inverse and matrix exponential.

### S3. Governing equations for the case of an axon with two equal branches (Fig. 1b)

#### S3.1. Equations expressing conservation of the total length of mitochondria in various compartments for an axon with two symmetric branches (Figs. 1b and 2b)

The 7^th^ demand site, simulating asymmetry between the branches, is removed. The model now consists of Eqs. (1)-(15). Eqs. (16)-(18), simulating transport in the 6^th^ demand site, are modified as follows (so that the 6^th^ demand site is symmetric with the 4^th^ demand site):

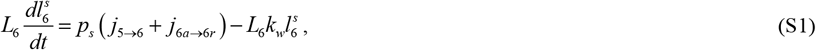

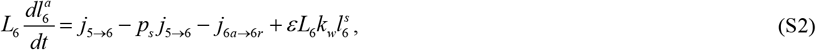

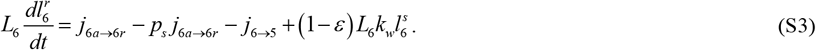

Mitochondria fluxes between the demand sites are simulated by Eqs. (22)-(25), (27)-(29), and (31)-(34). In addition, Eq. (35) is replaced with the following equation simulating the turn-around of mitochondria in the 6^th^ demand site:

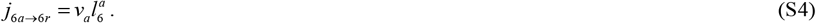

#### S3.2. Axon with two symmetric branches (Figs. 1b and 2b), changes in matrix B in Eq. (36)

For the case with two symmetric branches (Fig. 2b) matrix B is given by Eqs. (39)-(85), which are supplemented by the following equations:

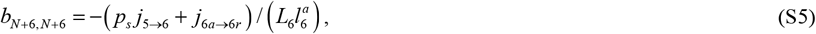

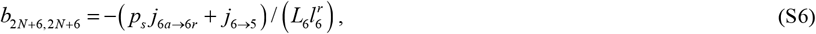

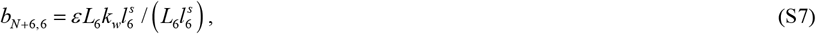

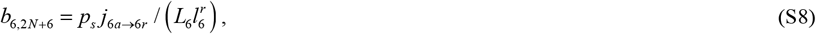

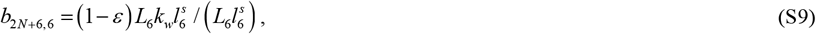

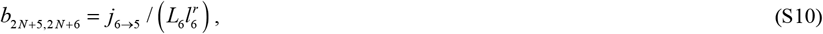

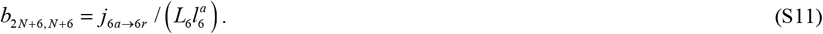

All other elements of matrix B, except for those given by Eqs. (39)-(85) and (S1)-(S11), are equal to zero.

## S4. Governing equations for the case of a straight axon (Fig. 1c)

### S4.1. Equations expressing conservation of the total length of mitochondria in various compartments for a straight (non-branched) axon (Figs. 1c and 2c)

Mitochondria concentrations in the first demand site in a straight axon (without branches) are simulated by Eqs. (1)-(3). Concentrations in the 2^nd^, 3^rd^, and 4^th^ demand sites are simulated by the following equations:

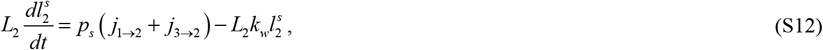

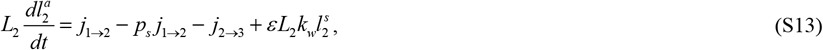

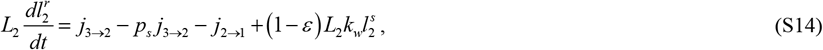

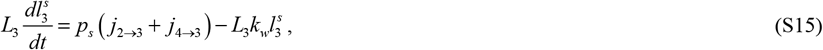

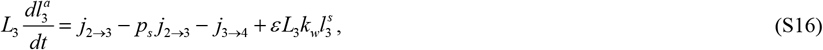

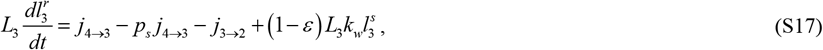

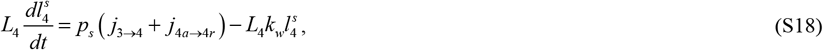

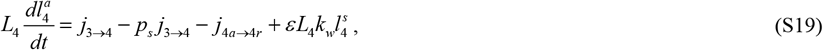

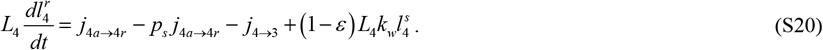

Fluxes between the compartments are given by Eqs. (22)-(24), (27)-(29), (33), and (34).

### S4.2. Straight (non-branched) axon (Figs. 1c and 2c), changes in matrix B in Eq. (36)

Elements of matrix B for the straight axon composed of four demand sites are given by Eqs. (39)-(49), which are supplemented by the following equations:

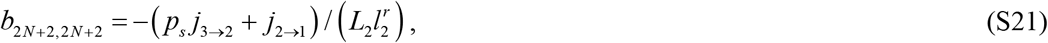

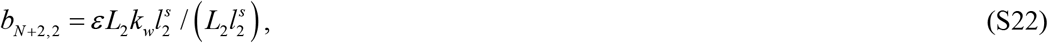

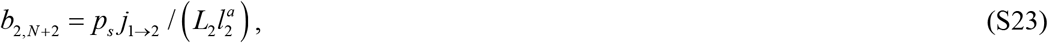

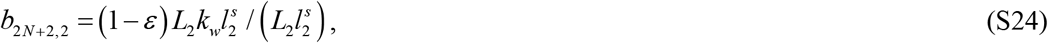

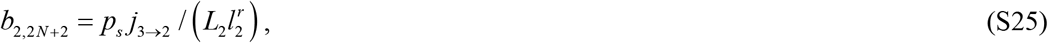

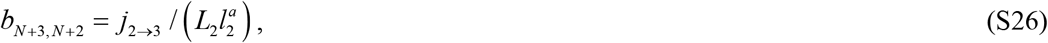

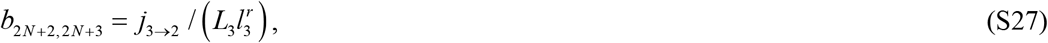

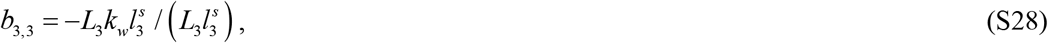

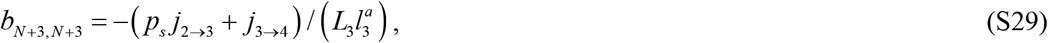

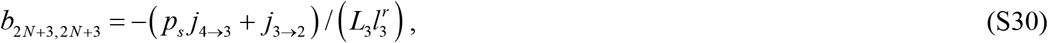

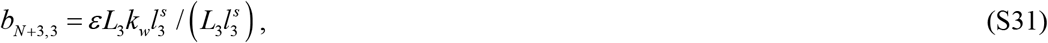

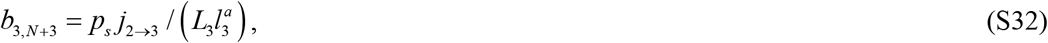

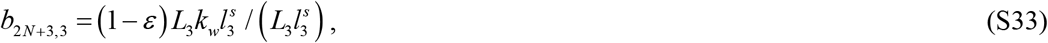

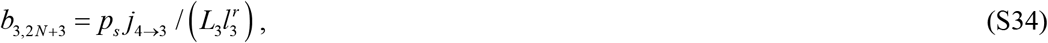

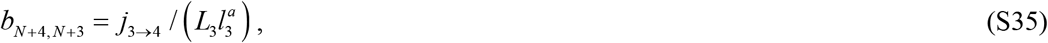

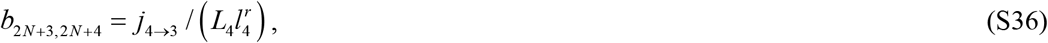

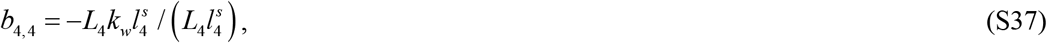

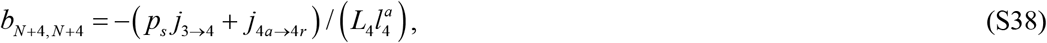

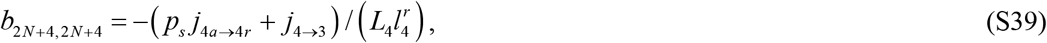

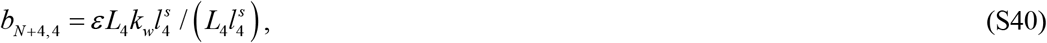

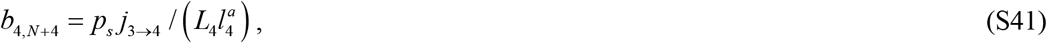

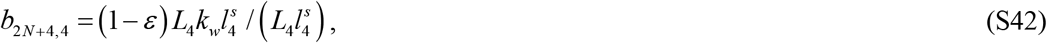

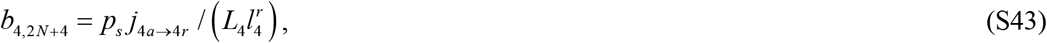

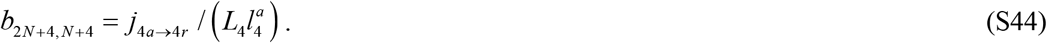

## S5. Supplementary figures

**Fig. S1.**
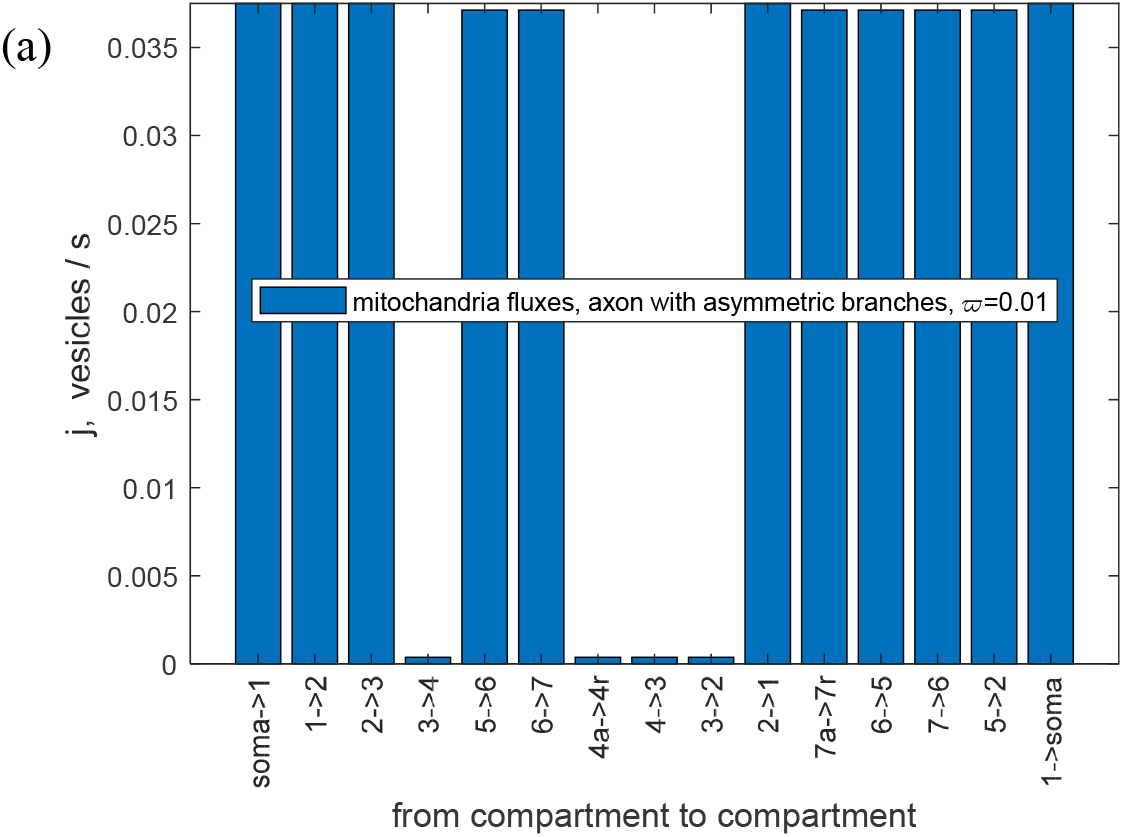
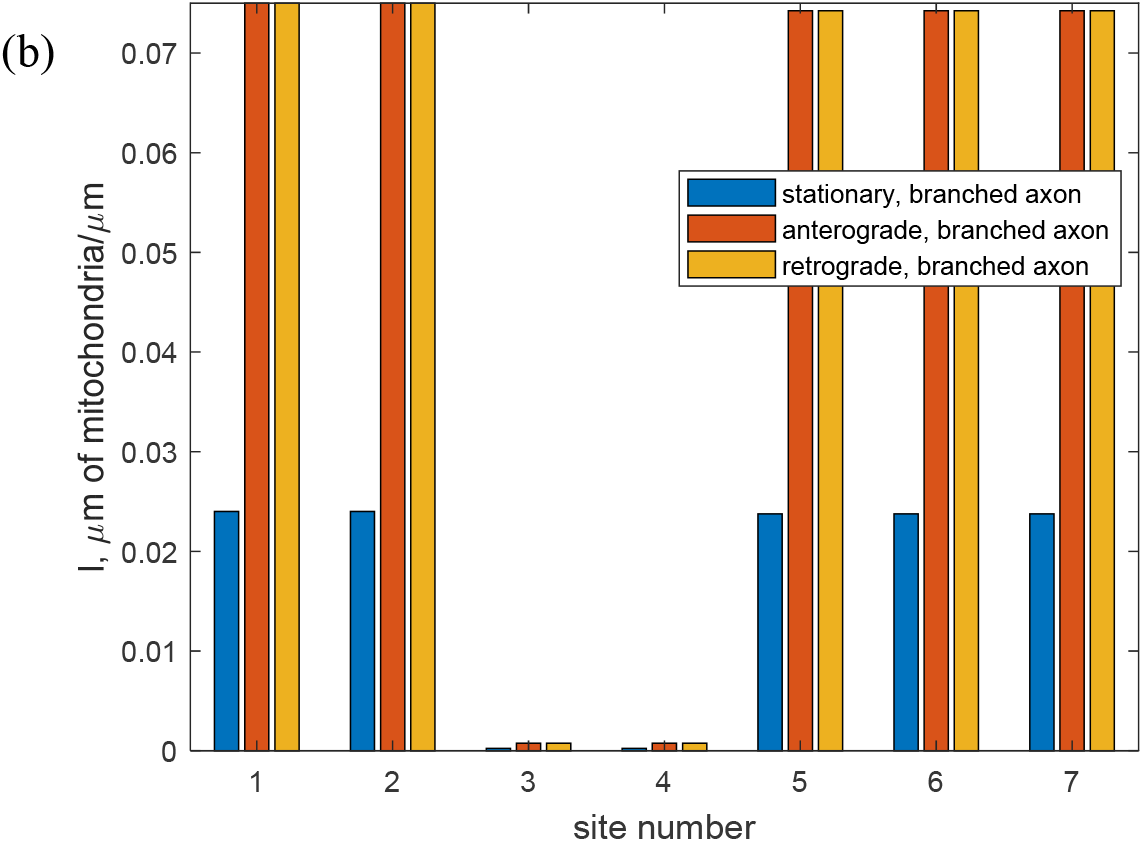
Branched axon with one shorter branch and one longer branch (Figs. 1a and 2a), steady state situation. (a) Fluxes of mitochondria traveling between the compartments. (b) Steady state values of the total length of stationary, anterogradely moving, and retrogradely moving mitochondria per unit length of the axon in the compartment by the *i*th demand site. *ϖ* = 0.01 (1% of mitochondria flux goes into the shorter branch and 99% goes into the longer branch).

**Fig. S2.**
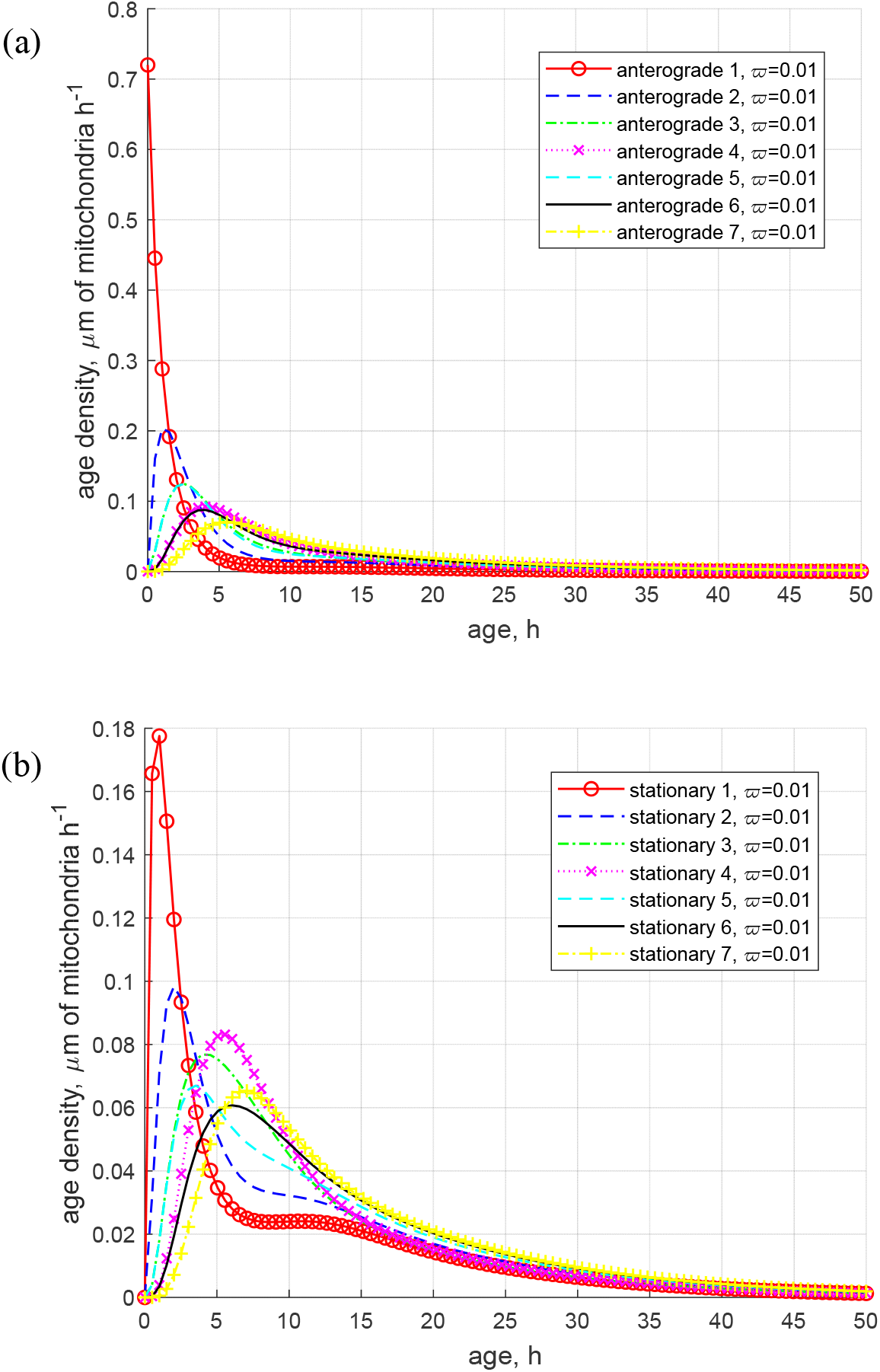
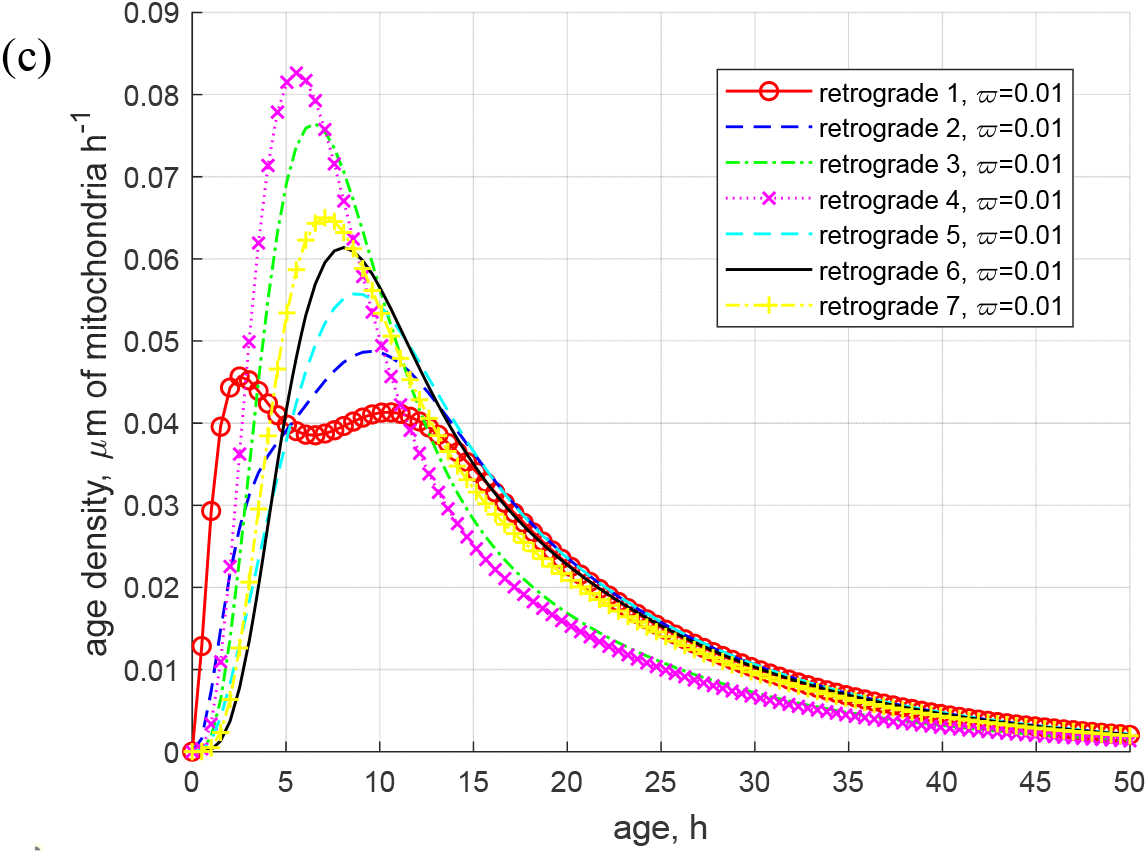
Branched axon with one shorter branch and one longer branch (Figs. 1a and 2a), steady state situation. (a) Age density of anterogradely moving mitochondria in various demand sites. (b) Age density of mitochondria in the stationary state in various demand sites. (c) Age density of retrogradely moving mitochondria in various demand sites. *ϖ* = 0.01 (1% of mitochondria flux goes into the shorter branch and 99% goes into the longer branch).

**Fig. S3.**
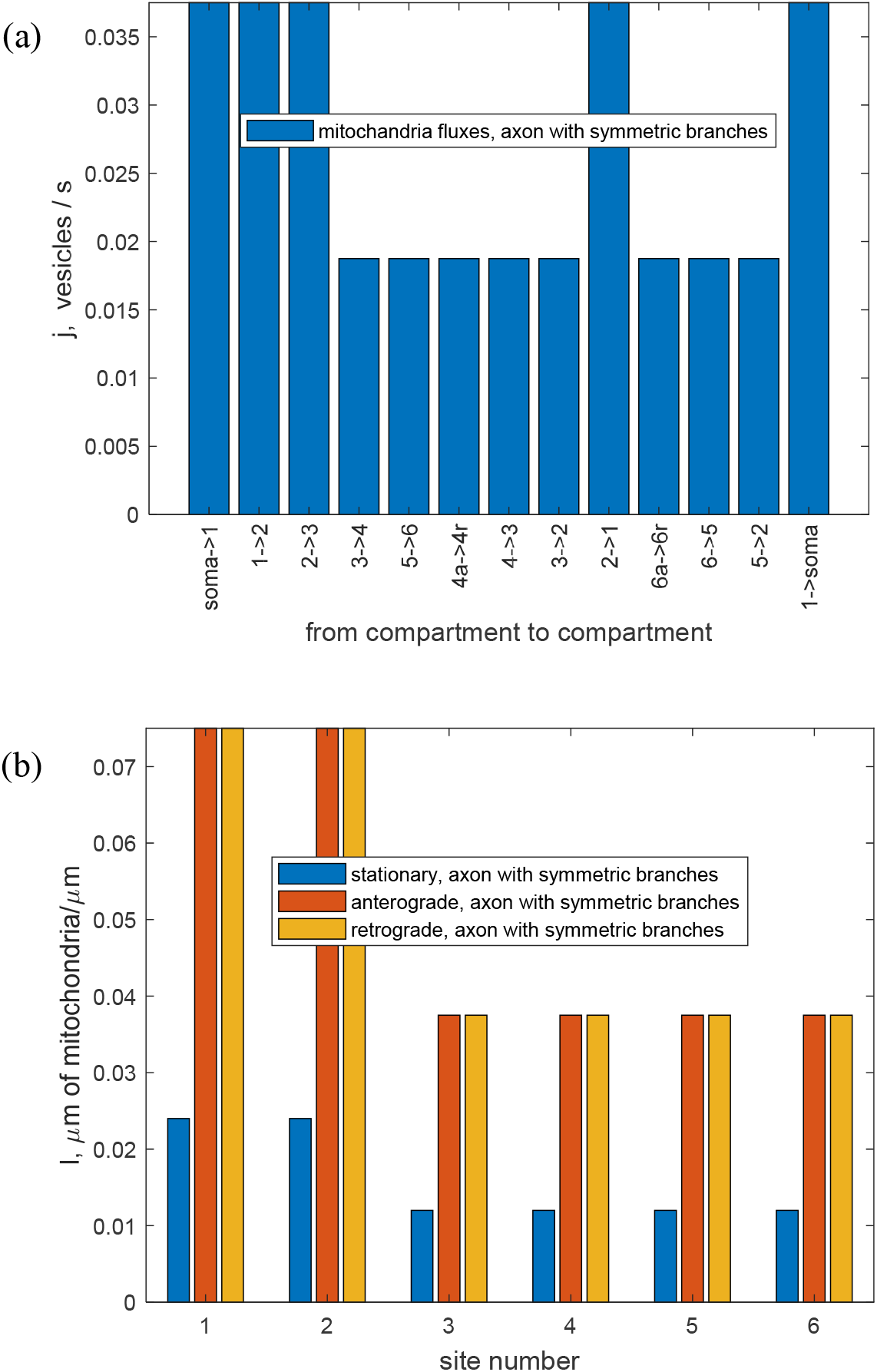
Branched axon with two equal branches (Figs. 1b and 2b), steady state situation. (a) Fluxes of mitochondria traveling between the compartments. (b) Steady state values of the total length of stationary, anterogradely moving, and retrogradely moving mitochondria per unit length of the axon in the compartment by the *i*th demand site. *ϖ* = 0.5 (the flux of mitochondria splits equally between the shorter and longer branches).

**Fig. S4.**
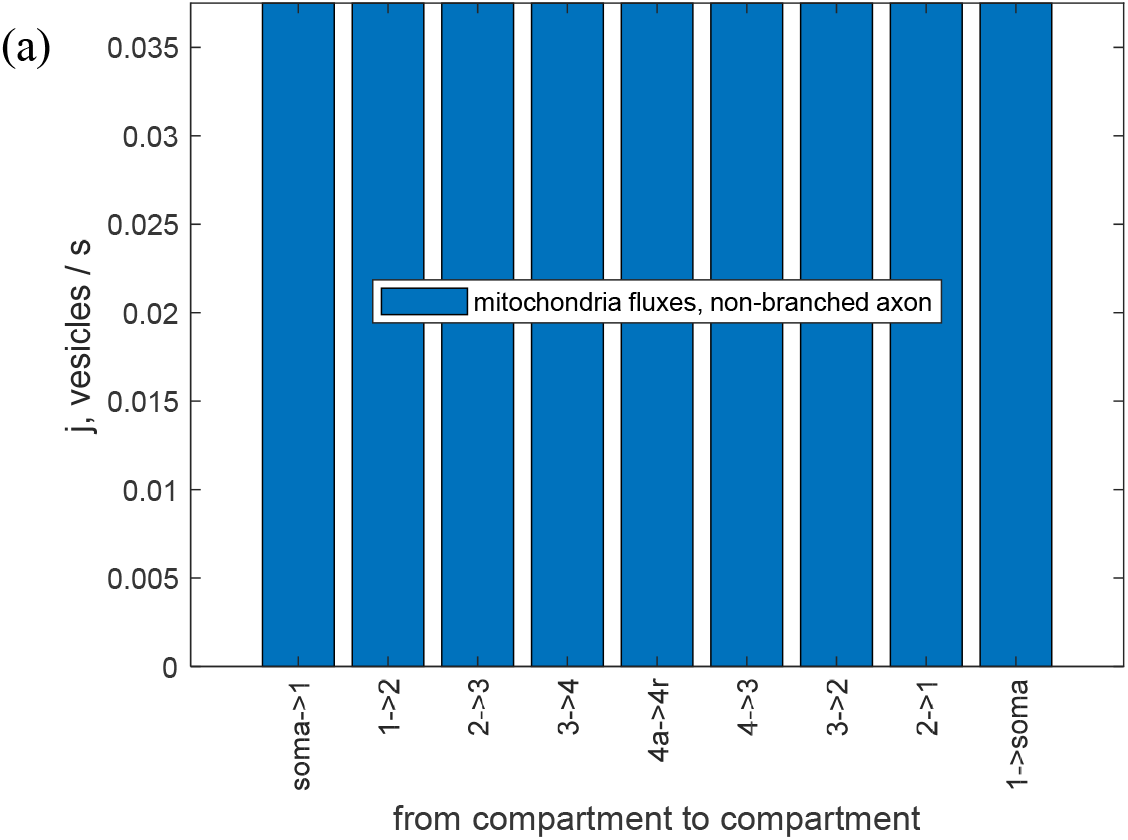
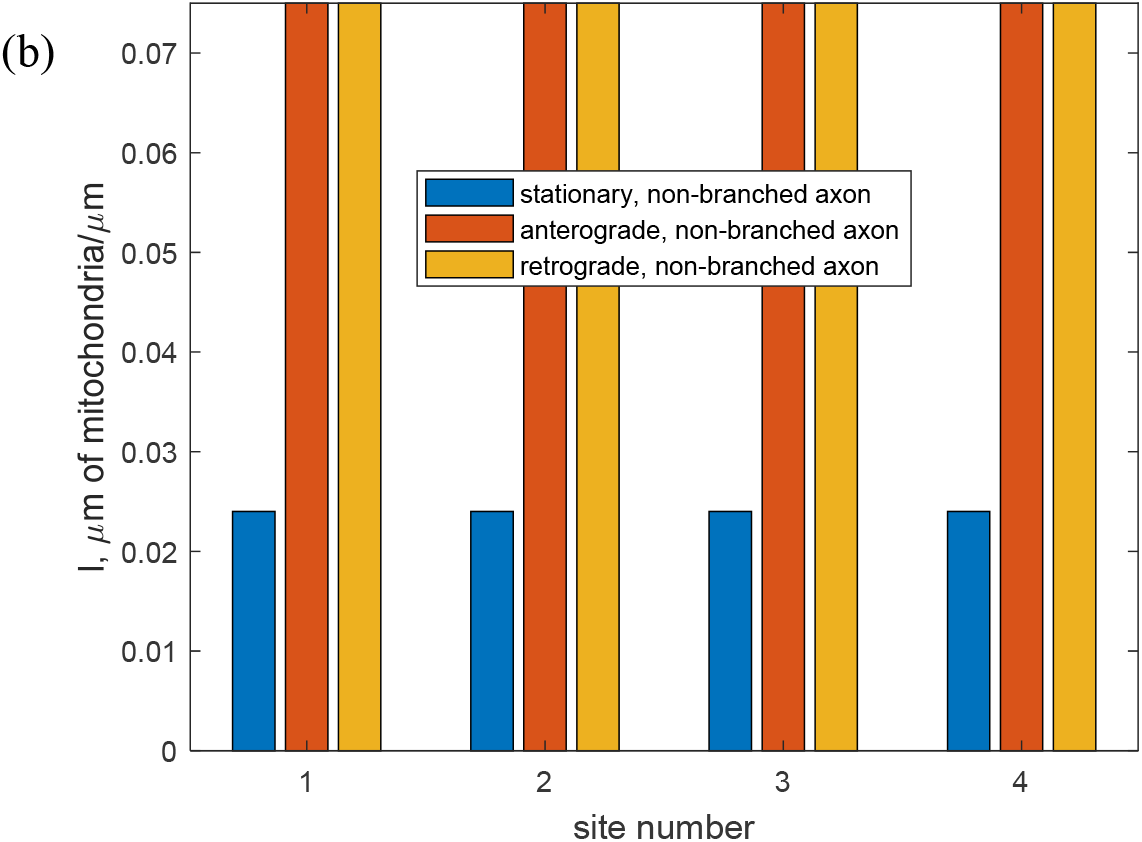
Non-branched (straight) axon (Figs. 1c and 2c), steady state situation. (a) Fluxes of mitochondria traveling between the compartments. (b) Steady state values of the total length of stationary, anterogradely moving, and retrogradely moving mitochondria per unit length of the axon in the compartment by the *i*th demand site.

